# Ubiquitin-based pathway acts inside chloroplasts to regulate photosynthesis

**DOI:** 10.1101/2022.06.06.494369

**Authors:** Yi Sun, Zujie Yao, Honglin Chen, Yiting Ye, Yuping Lyu, William Broad, Marjorie Fournier, Genyun Chen, Yonghong Hu, Shabaz Mohammed, Qihua Ling, R. Paul Jarvis

## Abstract

Photosynthesis is the energetic basis for most life on Earth, and in plants it operates inside double-membrane-bound organelles called chloroplasts. The photosynthetic apparatus comprises numerous proteins encoded by the nuclear and organellar genomes. Maintenance of this apparatus requires the action of internal chloroplast proteases, but a role for the nucleocytosolic ubiquitin-proteasome system (UPS) was not expected owing to the barrier presented by the double-membrane envelope. Here, we show that photosynthesis proteins (including those encoded internally by chloroplast genes) are ubiquitinated, and processed via the CHLORAD pathway: they are degraded by the 26S proteasome following CDC48-dependent retrotranslocation to the cytosol. This demonstrates that the reach of the UPS extends to the interior of endosymbiotically-derived chloroplasts, where it acts to regulate one of the most fundamental processes of life.

**One Sentence Summary:** The ubiquitin-proteasome system targets proteins inside chloroplasts, including chloroplast-encoded photosynthesis subunits.

## Main text

Chloroplasts are the defining organelles of plants and algae, and evolved through endosymbiosis from a cyanobacterial ancestor (*1*). They use light energy to convert CO_2_ into carbohydrate through photosynthesis, and thus play a pivotal role in controlling atmospheric CO_2_ levels. Apart from photosynthesis, chloroplasts have a multiplicity of metabolic functions (including the synthesis of amino acids, fatty acids, and plant hormones), and possess a correspondingly diverse proteome (*2*). Homeostatic mechanisms governing the chloroplast proteome, including proteolysis, are consequently of vital importance for organellar functions and plant development.

The chloroplast proteome comprises approximately 3000 proteins, >90% of which are encoded in the nucleus with the remainder encoded by the chloroplast genome (*1, 2*). Several classes of internal protease of prokaryotic origin participate in chloroplast proteostasis, and many of these contribute to the regulation of photosynthesis (*3, 4*). Photosynthesis is a highly complex process carried out by distinct functional units, each one with numerous protein components: photosystems I and II (PSI, PSII), linked by the cytochrome b_6_f complex, for light harvesting; ATP synthase; and the Calvin cycle for carbon fixation. Proteolytic regulation of the D1 subunit of the PSII reaction centre by internal FtsH and Deg proteases, which is crucial for maintaining photosynthetic efficiency, has been studied intensively (*5-7*). However, mechanisms underlying the proteolysis of many other photosynthesis components remain unclear.

Our recent work uncovered a role for the ubiquitin-proteasome system (UPS) in the regulation of chloroplast outer envelope membrane (OEM) proteins, via a pathway termed CHLORAD (for chloroplast-associated protein degradation) (*8, 9*). CHLORAD involves the RING (really interesting new gene)-type ubiquitin E3 ligase SP1, the Omp85-type β-barrel channel SP2, and the AAA+ (ATPase associated with diverse cellular activities) chaperone CDC48. The SP1 and SP2 proteins form a complex in the OEM, with SP1 mediating ubiquitination of target proteins and SP2 forming a retrotranslocon for the delivery of ubiquitinated targets to the cytosol, for degradation by the 26S proteasome. The motive force for such retrotranslocation is provided by CDC48. Known targets of CHLORAD are the TOC proteins that mediate chloroplast protein import (*10-13*), the regulation of which allows CHLORAD to control organellar development and functions, for example in response to stress conditions (*14*).

In contrast with the functionally analogous ERAD system which eliminates a broad range of misfolded and functional ER proteins (*15, 16*), the data published to-date suggest that CHLORAD has a narrower focus on the protein import machinery in the chloroplast OEM. Given that CHLORAD is heavily involved in the profound structural and functional changes that chloroplasts (and other plastid types) undergo during plant development (*8, 9, 13, 17*), it is conceivable that CHLORAD has a broader role that includes, for example, the removal of those proteins that become redundant during such dynamic processes. As chloroplasts have a diversity of functions (*1, 2*), it is possible that many as yet unknown substrates of CHLORAD exist in chloroplasts, particularly in the interior which accounts for more than 99% of total chloroplast protein (*18*).

## Results

### Detection of internal chloroplast ubiquitination by fractionation and affinity purification

To gain initial insight into the possibility that CHLORAD is wider in scope than previously envisaged, we conducted an immunoblot investigation of isolated *Arabidopsis* chloroplasts. To improve the sensitivity of this analysis, a transgenic plant line expressing Myc-tagged ubiquitin (6Myc-Ub) was employed. We detected high-molecular-weight smears in purified chloroplasts upon anti-Myc analysis (Fig. 1A). Moreover, these anti-Myc smears persisted following treatment of the chloroplasts with thermolysin (a protease that removes only surface-exposed OEM proteins) (*19*) (Fig. 1B), indicating that ubiquitinated proteins may exist in internal compartments, such as the inner envelope membrane (IEM), stroma, or thylakoid membranes. Support for the existence of ubiquitinated proteins in the stroma (the main aqueous compartment of the organelle) was provided by subfractionation, which revealed the presence of polyubiquitin smears in both membrane and soluble (predominantly stroma) fractions (Fig. 1C).

**Fig. 1.**
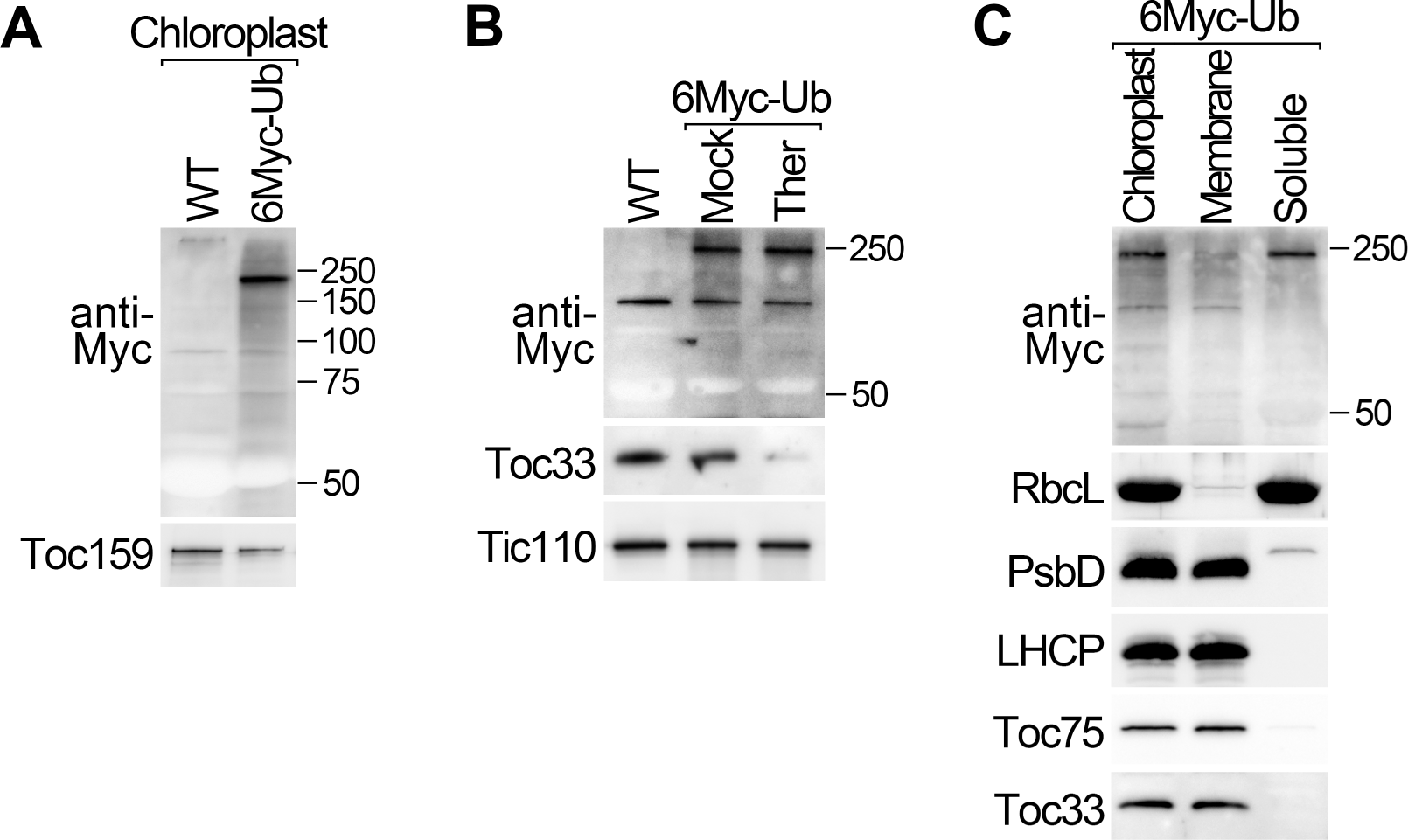
Proteins of the chloroplast interior are polyubiquitinated. (**A** and **B**) Chloroplasts isolated from transgenic plants expressing Myc-tagged ubiquitin (6Myc-Ub), and from wild type, were analysed by immunoblotting (A). Similar chloroplasts were treated with thermolysin protease (Ther), or buffer lacking protease (Mock), before immunoblotting analysis (B). The asterisk in B indicates a non-specific band. (**C**) Chloroplasts isolated from 6Myc-Ub plants were separated into membrane pellet (P18) and soluble supernatant (S18; predominantly stroma) fractions by centrifugation at 18,000 × *g*, and then analysed by immunoblotting. Analysis of control proteins confirmed the efficacy of the protease treatment and fractionation steps. Positions of molecular weight markers (sizes in kD) are shown to the right of the images.

To verify that the observed smears indeed correspond to ubiquitinated chloroplast proteins, they were enriched by anti-Myc immunoprecipitation and analysed by mass spectrometry. A number of internal chloroplast proteins were identified, and for one of them a specific ubiquitination site was detected: this was the stromal protein PrfB3, which functions in chloroplast gene expression (fig. S1, table S1) (*20*). Thus, altogether, these data supported the hypothesis that CHLORAD acts on proteins inside chloroplasts.

### Ubiquitinomics reveals that proteins involved in photosynthesis are ubiquitinated

To develop a more complete picture of the substrates of CHLORAD, we sought to systematically identify ubiquitination targets in chloroplasts, and their sites of modification. First, chloroplasts purified from wild-type plants were analysed using a ubiquitin remnant (di-Gly) antibody-based peptide enrichment ubiquitinomic approach (*21*), and the samples were subjected to proteomic analysis. In total, 57 non-redundant ubiquitination sites in 40 proteins were identified (table S2). To increase the sensitivity of detection, retrotranslocation of CHLORAD substrates was blocked using a dominant-negative mutant of CDC48 (CDC48-DN) (*9*); indeed, a much stronger ubiquitination signal could be detected in isolated chloroplasts following CDC48-DN induction (fig. S2A). Chloroplasts purified from CDC48-DN plants were analysed using the same approach (*21*), and three independent experiments were performed. The ubiquitinated proteins were identified by cross-referencing with a custom-assembled chloroplast proteome database comprising 4174 proteins (*18, 22, 23*). In total, 768 non-redundant ubiquitination sites in 316 proteins were identified in at least one experiment for CDC48-DN (table S3, fig. S2). These results indicated that the chloroplast proteome is broadly ubiquitinated, and that ubiquitinated chloroplast proteins are commonly processed by CDC48.

The identified putative CHLORAD substrates belonged to various functional categories, but with a noteworthy enrichment in photosynthesis components (Fig. 2A). With regard to suborganellar localization, OEM proteins were well represented in the set (20 out of 316 proteins), which is consistent with our published results (*8, 9*); TOC components (Toc159, Toc75 and Toc34) and SP1 and SPL2 were present as expected, as well as others such as fatty acid (FA) synthetase LACS9 and the channel protein OEP24 (*18, 24*). However, a considerable number of IEM, stromal and thylakoidal proteins were also identified, including the lipoxygenase LOX2 (*25*), Rubisco small subunit RbcS, various metabolism-related proteins, light harvesting chlorophyll-binding proteins, and photosystem subunits such as PsaA, PsaB and PsbC (*26*) (Fig. 2B, table S3). This suggested that ubiquitination is a major mechanism for the regulation of the chloroplast proteome, particularly regarding photosynthesis.

**Fig. 2.**
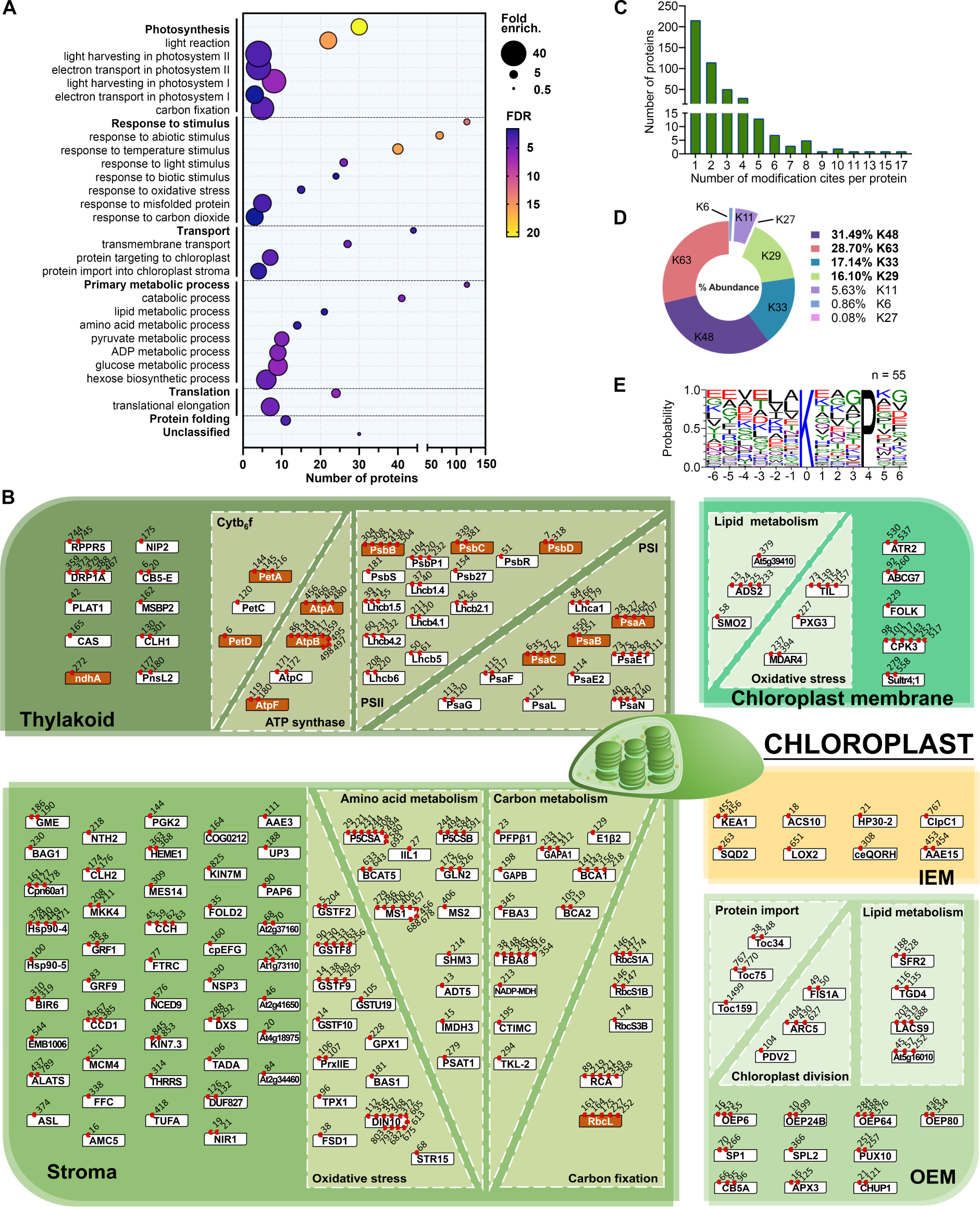
Photosynthesis and other proteins of the chloroplast interior are prominent in the chloroplast ubiquitinome. (**A**) Dot plot showing significantly overrepresented GO terms in the chloroplast ubiquitinome, as determined using chloroplasts purified from CDC48-DN plants after estradiol induction. Dot size indicates overrepresentation (fold enrichment) compared to the whole genome. Dot colour indicates False Discovery Rate (FDR; -log_10_ [*P* value]), where higher FDR values indicate more statistically significant enrichment. Dots are not shown for terms lacking statistically significant (*P* < 0.05) enrichment. (**B**) Suborganellar and functional distribution of the chloroplast ubiquitinome, showing also ubiquitination sites as determined by diGly analysis. Localizations were assigned manually and only proteins localized in a defined chloroplast compartment (OEM, IEM, stroma and thylakoid), or in an internal chloroplast membrane fraction (IEM or thylakoid), are shown. Boxes indicate individual proteins (white, nucleus-encoded; orange, chloroplast-encoded), and red circles with numbers indicate which amino acids showed ubiquitination. (**C**) Histogram showing the number of ubiquitination sites detected per protein in the chloroplast ubiquitinome. (**D**) Pie chart showing the relative abundance (based on peptide intensity) of different polyubiquitin linkage types in the chloroplast ubiquitinome. Values are means from three experiments. (**E**) Logo plot showing motif analysis of ubiquitinated peptides in the chloroplast ubiquitinome. Six residues either side of the modification site (position 0) are shown.

Remarkably, among the putative CHLORAD substrates were 13 chloroplast-encoded proteins with identified ubiquitination sites. These included two PSI subunits, four PSII subunits, two cytochrome b_6_f subunits, three ATP synthase subunits, one NDH complex subunit, and the Rubisco large subunit (table S4). This was an important finding as it ruled out the possibility that the detected ubiquitination was the consequence of modification of unimported cytosolic precursors (preproteins) (*27*); although we note that this was already highly unlikely given that the ubiquitinome analysis was conducted using purified chloroplasts.

To verify the results of the ubiquitinome analysis, selected proteins were analysed using an in vivo ubiquitination assay. Cells expressing FLAG-tagged ubiquitin (FLAG-Ub) were subjected to anti-FLAG immunoprecipitation, and the eluates were probed using antibodies against the putative CHLORAD substrates. This confirmed the ubiquitination of the stromal PrfB3 protein (*20*) (fig. S3A), as well as that of proteins encoded by the chloroplast genome (i.e., PsaA and PsbC) (fig. S3B). Moreover, the amount of detectable ubiquitination was enhanced when substrate retrotranslocation was blocked by expressing CDC48-DN (fig. S3B). This provided further strong support for the conclusion that proteins of the chloroplast interior are regulated by ubiquitination and CDC48.

The chloroplast ubiquitinome had 2.7 ubiquitination sites per protein, suggesting extensive ubiquitination of chloroplast proteins (Fig. 2C); the global ubiquitinome has 1.1 sites per protein (*28*). Analysis of the di-Gly data revealed that all polyubiquitin linkage types are present in chloroplasts, with the following distribution, based on integration of signal: Lys-48 > Lys-63 > Lys-33 > Lys-29 > Lys11 >>> Lys-6/Lys-27 (Fig. 2D). Because K48 was the most abundant linkage type, much of chloroplast ubiquitinome can be assumed to be for proteasomal degradation. Intriguingly, we identified a putative consensus motif for ubiquitin attachment (Fig. 2E), which was not observed previously in the global plant ubiquitinome (*28-30*), implying that a specific ubiquitination process may occur inside chloroplasts. In addition, ubiquitination sites were identified in different sub-chloroplastic compartments, although the transmembrane domains clearly lack ubiquitination (fig. S4).

### Quantitative proteomics reveals that CHLORAD regulates the levels of a broad range of proteins

A hallmark of CHLORAD substrates is that their steady-state levels increase upon CHLORAD inhibition. Thus, as a complement to the ubiquitinome analysis, we sought chloroplast proteins that over-accumulate after CDC48 inhibition. To this end, we performed label-free quantitative proteomic analysis of plants expressing CDC48-DN or the corresponding CDC48-WT control, and then filtered the data using the same custom chloroplast proteome database as used for the ubiquitinomics; this approach avoided the need for a time-consuming chloroplast isolation step which might bias protein quantification. In three biological replicates, we detected 1444 proteins in both genotypes of which ∼27% were present at elevated levels (>1.5 fold) in CDC48-DN relative to the control, as expected for CHLORAD targets (Fig. 3A and B, fig. S5, table S5); a selection of these data are shown in table S6. The results were highly consistent with our published immunoblot data (*9*): TOC components (Toc159, Toc33 and Toc75) were over-accumulated in CDC48-DN, whereas established non-substrate proteins (Tic40 and OEP80) were unchanged (table S6). We also conducted label-free quantitative proteomic analysis of chloroplasts isolated from the same CDC48-DN and CDC48-WT plants, and similar results were obtained (table S7).

**Fig. 3.**
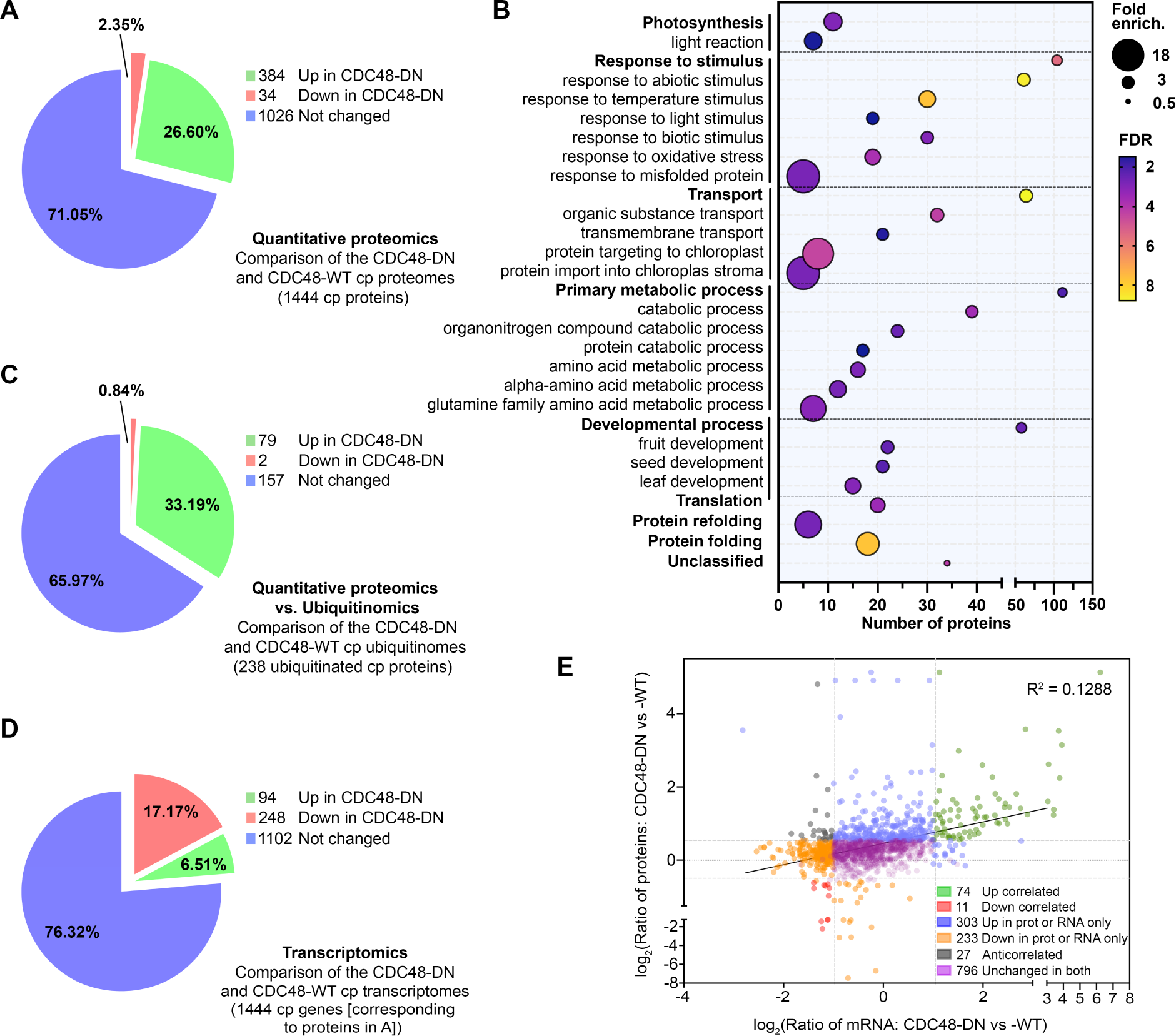
Identification of numerous CHLORAD substrates using quantitative proteomics. (**A**) Pie chart showing the differential accumulation of chloroplast (cp) proteins upon CHLORAD inhibition, as determined by quantitative proteomic analysis of CDC48-DN and CDC48-WT transgenic plants after estradiol induction. (**B**) Dot plot showing significantly overrepresented GO terms for chloroplast proteins that are over-accumulated in CDC48-DN plants. Dot size indicates overrepresentation (fold enrichment) compared to the whole genome. Dot colour indicates False Discovery Rate (FDR; -log_10_ [*P* value]), where higher FDR values indicate more statistically significant enrichment. Dots are not shown for terms lacking statistically significant (*P* < 0.05) enrichment. (**C**) Pie chart showing the differential accumulation of ubiquitinated chloroplast proteins in the quantitative proteomic analysis depicted in A. Only proteins in the chloroplast ubiquitinome (Fig. 2) are shown. (**D**) Pie chart showing the differential expression of mRNAs corresponding to the chloroplast proteins identified by proteomics in A, as determined by RNA-seq transcriptomics. (**E**) Dot plot showing a lack of correlation between chloroplast protein abundancies (from A) and corresponding mRNA abundancies (from D). The coefficient of determination (R^2^) value is shown.

Among the other elevated proteins in CDC48-DN were a number of OEM proteins unrelated to the core TOC apparatus, including LACS9, OEP64, OEP24 (a channel protein), and CHUP1 (which regulates chloroplast movement), all of which were identified also by ubiquitinomics (*18, 24, 31*). Given that the OEM proteome is much less abundant than those of the stroma and thylakoids (*22*), this suggested that OEM proteins are major substrates of CHLORAD, as would be expected. In a further parallel with the ubiquitinome analysis, many IEM and stromal proteins (e.g., LOX2) were also elevated in CDC48-DN. These internal proteins have a range of different functions, including photosynthesis, chlorophyll metabolism, and oxidative stress response.

To verify the proteomics data, immunoblotting and confocal microscopy analyses were performed. Some proteins were analysed using specific antibodies, and their levels were found to be elevated in CDC48-DN plants, as expected (fig. S6). Where specific antibodies were not available, selected proteins (including LACS9 and FAX1) were transiently expressed with a YFP tag, and then analysed by immunoblotting and microscopy (fig. S7). Both analytical approaches showed the proteins to be elevated in CDC48-DN cells, while the latter revealed that the accumulated protein is localized to the chloroplasts in each case (fig. S5). Notably, of the 238 proteins detected in this proteomic analysis that also contain at least one ubiquitination site (tables S3 and S5), 79 (33%) were over-accumulated in CDC48-DN chloroplasts (Fig. 3C), supporting the notion that they are CHLORAD substrates.

Moreover, RNA sequencing (RNA-seq) analysis revealed that, although there are many significant CDC48-dependent changes in global transcription (fig. S8, table S8), including of ubiquitin-related proteolytic genes, most transcripts encoding proteins over-accumulated in CDC48-DN chloroplasts were not up-regulated transcriptionally (Fig. 3D and E). In fact, mRNAs for chloroplast proteins tended to be reduced in CDC48-DN plants, in contrast to the protein levels (Fig. 3D). In general, we found no correlation between mRNA and protein levels for the chloroplast proteins identified in our quantitative proteomics analysis, implying that the protein abundance changes are mediated post-translationally.

Overall, based on the ubiquitinomic and quantitative proteomic analyses, we concluded that CHLORAD functions much more broadly than was previously envisaged, by acting on many proteins of the chloroplast’s interior.

### Demonstrating CHLORAD involvement in the degradation of internal chloroplast proteins

Identification of many internal chloroplast proteins (non-OEM proteins) as putative CHLORAD substrates was surprising, as such proteins are separated from the cytosolic UPS apparatus by the envelope membranes. To confirm that these proteins are indeed processed by the UPS, we examined their degradation following proteasome inhibition. We began with the nucleus-encoded stromal protein PrfB3. We monitored levels of a PrfB3-HA fusion protein following inhibition of protein synthesis using cycloheximide, and found that degradation was apparent within 6 hours, indicating brisk turnover. Crucially, degradation of PrfB3-HA was delayed following treatment with bortezomib which suppresses proteasome and CHLORAD activity (Fig. 4A and B) (*9*).

**Fig. 4.**
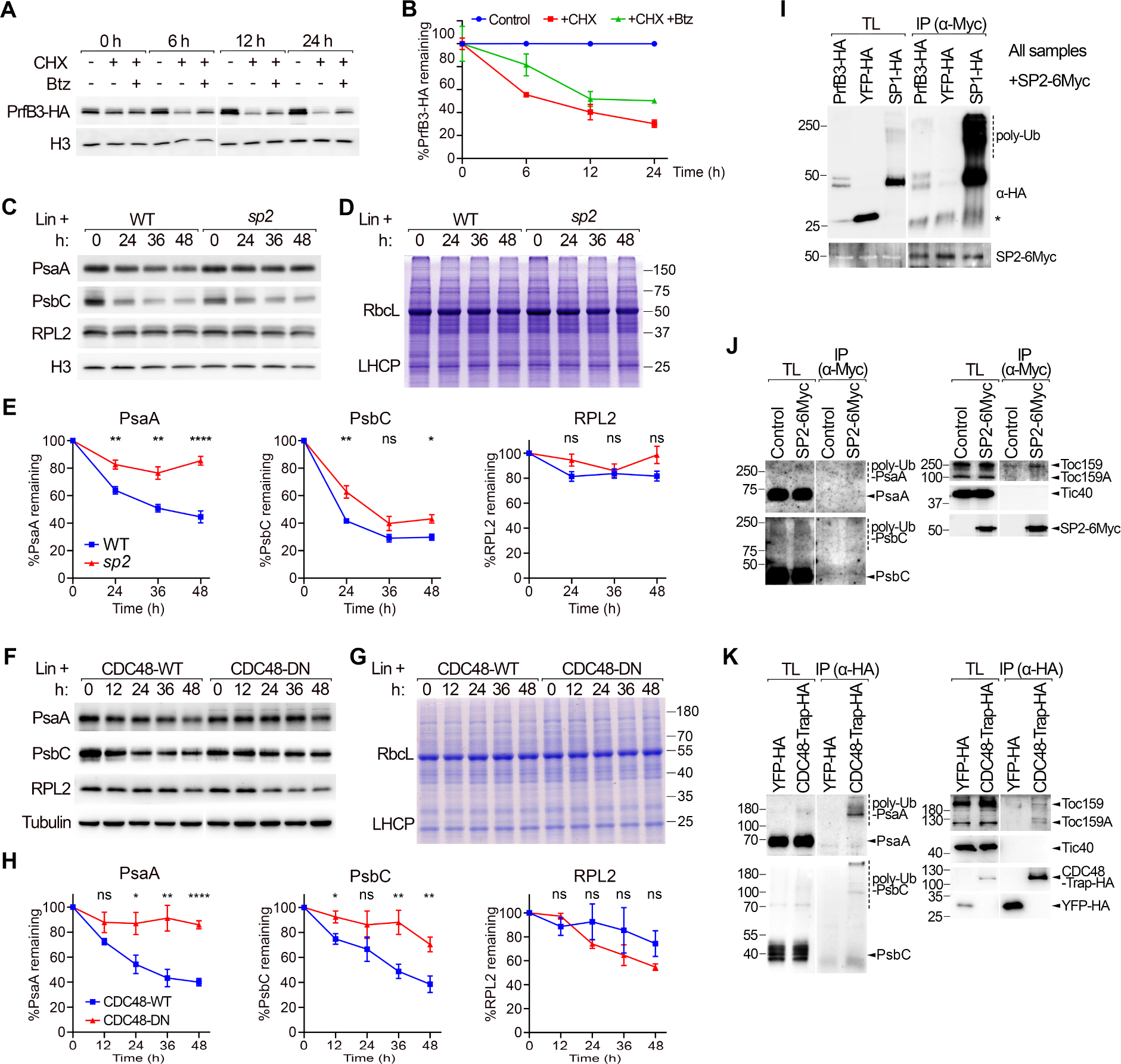
Proteins of the chloroplast interior are processed by CHLORAD. (**A** and **B**) Proteasome dependency of the degradation of nucleus-encoded PrfB3 was analysed after inhibiting translation using cycloheximide (CHX). Wild-type protoplasts expressing PrfB3-HA were incubated with or without CHX and the proteasome inhibitor bortezomib (Btz) for the indicated times, and then analysed by immunoblotting (A). Band intensities for PrfB3-HA were quantified and normalized to equivalent data for a histone H3 loading control (B). Time 0 was taken as 100%. Data are means ± SEM from at least two experiments. (**C** to **E**) SP2 dependency of the degradation of chloroplast-encoded proteins was analysed after inhibiting translation using lincomycin (Lin). Wild-type and *sp2* mutant plants were incubated with Lin for the indicated times, and then analysed by immunoblotting (C). Coomassie staining indicated that the treatments did not change the overall protein profile (D). Band intensities were quantified and normalized to equivalent histone H3 data (E). Time 0 was taken as 100%. Data are means ± SEM from at least four experiments. Asterisks in E indicate significance according to an unpaired two-tailed Student’s t-test (*, significant at *P* < 0.05; **, significant at *P* < 0.01; ****, significant at *P* < 0.0001; ns, no statistical significance). (**F** to **H**) CDC48 dependency of the degradation of chloroplast-encoded proteins was analysed after inhibiting translation using Lin. Data are equivalent to those in C-E, except that CDC48-WT and CDC48-DN transgenic plants were used. Statistical analysis in H is as defined for E. (**I** to **K**) Interaction of putative substrates with the CHLORAD apparatus was assessed by co-immunoprecipitation (IP). In I, wild-type protoplasts were co-transfected with the *SP2-6Myc* construct and either *PrfB3-HA*, *YFP-HA* or *SP1-HA*. In J, wild-type or SP2-6Myc seedlings were analysed without protoplastation. In K, wild-type protoplasts were transfected with the *YFP-HA* or *CDC48-Trap-HA* (AtCDC48A^E581Q^; which displays stabilized substrate binding (*9*)) constructs. Immunoprecipitations were performed using anti-Myc (I,J) or anti-HA (K) resin, and analysed by immunoblotting using the indicated antibodies. In all cases, YFP-HA acted as a negative control. Positions of molecular weight markers (sizes in kD) are shown to the left of the images. The asterisk in I indicates a non-specific band. TL, total lysate; poly-Ub, poly-ubiquitinated forms.

Next, we analysed the turnover of several chloroplast-encoded proteins in the CHLORAD-defective *sp2* and CDC48-DN backgrounds (*9*). In this case, native protein levels were monitored following treatment with lincomycin to inhibit chloroplast translation. The stabilities of PsaA and PsbC (which are both putative substrates, and photosynthesis components) were clearly enhanced in the CHLORAD-defective genotypes (Fig. 4C to H). In contrast, RPL2 (a chloroplast-encoded ribosomal protein not identified as a putative CHLORAD substrate, used here as a control) was not stabilized in the *sp2* and CDC48-DN backgrounds. Thus, CHLORAD acts selectively on a diverse range of internal substrates, but not all chloroplast proteins.

To further establish a direct link between the CHLORAD apparatus and its putative targets in the chloroplast interior, we performed co-immunoprecipitation experiments. SP2-6Myc was found to specifically associate with PrfB3-HA, including high molecular weight forms that we interpret to be polyubiquitinated (Fig. 4I). Similarly, SP2-6Myc and a CDC48 mutant with stabilized substrate binding (CDC48-Trap) (*9*) associated with two of the putative chloroplast-encoded CHLORAD substrates (PsaA and PsbC) (Fig. 4J and K). Therefore, the action of CHLORAD on the stability of the internal substrates may be mediated through direct physical interactions.

### Demonstrating the role of CDC48 in the extraction of internal chloroplast proteins

We previously demonstrated that CDC48 drives the retrotranslocation of CHLORAD substrates from the OEM to the cytosol. To assess whether CDC48 is similarly involved in the extraction of substrates from the chloroplast interior, we performed in vivo retrotranslocation assays for two chloroplast-encoded substrates (PsaA and PsbC) using our established system (*9*). These substrates were selected to eliminate the possibly of confounding effects due to the accumulation of unimported preproteins in the cytosol.

Accordingly, we separated bortezomib-treated protoplasts from CDC48-DN and CDC48-WT plants into chloroplast and cytosol fractions, and then enriched the ubiquitinated proteins from each fraction by immunoprecipitation. Resident and extracted PsaA and PsbC proteins, in the chloroplast and cytosol samples respectively, were then detected by immunoblotting (Fig. 5A). The extraction of polyubiquitinated PsaA and PsbC to the cytosol was clearly apparent, providing strong evidence that internal chloroplast protein can be exported from the organelle. Moreover, such extraction was inhibited in cells expressing CDC48-DN, as indicated by the lack of ubiquitinated material in the cytosolic fractions, pointing to an important role for CDC48 in the processing of such internal chloroplast proteins (Fig. 5).

**Fig. 5.**
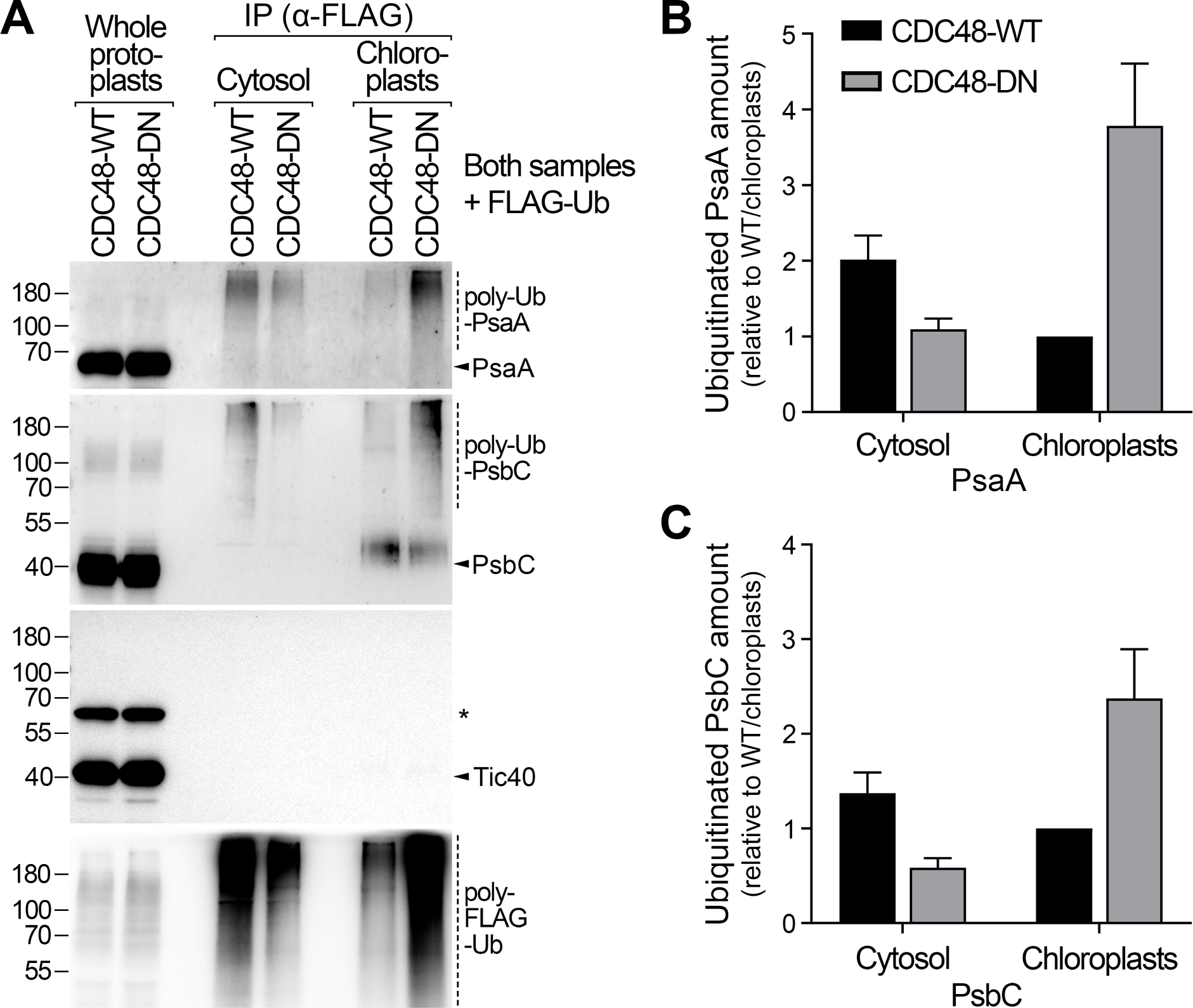
CDC48 is required for the extraction of polyubiquitinated photosynthesis proteins. (**A**) The role of CDC48 in the extraction of chloroplast-encoded PsaA and PsbC substrates was analysed using an in vivo retrotranslocation assay. Protoplasts from CDC48-WT and CDC48-DN transgenic plants that were transiently expressing FLAG-Ub were treated, after estradiol induction, with bortezomib proteasome inhibitor and then separated into cytosol and chloroplast fractions. In this assay, retrotranslocation occurred in intact cells, and the extracted substrates were protected by bortezomib inhibition, which initiated the experiment. After fractionation, ubiquitinated proteins were immunoprecipitated from both fractions and detected by immunoblotting. Tic40, which is not a substrate of CHLORAD, served as a negative control. Typical immunoblotting results are shown. Positions of molecular weight markers (sizes in kD) are shown to the left of the images. The asterisk indicates a non-specific band. (**B** and **C**) Retrotranslocation efficiency was assessed by quantifying the relative amounts of ubiquitinated PsaA (B) and PsbC (C) in the cytosol and chloroplast fractions described in A. Data are means ± SEM from at least three experiments.

### Evaluating the physiological importance of CHLORAD for photosynthesis and lipid homeostasis

Short-term expression of CDC48-DN causes chlorosis (Fig. 6A) (*9*), which was previously linked to over-accumulation of reactive oxygen species (ROS) due to a failure to properly regulate chloroplast protein import (*9*). Such chlorosis also suggested morphological changes in the chloroplasts. To investigate the latter possibility, we assessed chloroplast ultrastructure in these plants. Chloroplasts in CDC48-DN plants contained enlarged plastoglobules (Fig. 6B and C), which is consistent with oxidative stress (*32*) caused by disrupted photosystem component homeostasis, and were smaller than wild-type chloroplasts (Fig. 6B and D), which can explain the chlorotic phenotype of the seedlings. Moreover, CDC48-DN chloroplasts contained larger grana (stacked thylakoids, where PSII is concentrated (34)) and fewer stromal thylakoids (Fig. 6B and E). These observations, alongside the data showing that CHLORAD acts selectively on a range of chloroplast-encoded photosystem components, suggested that CHLORAD plays a nuanced role in regulating the activities of the photosystems. To address this hypothesis, we simultaneously measured energy conversion in PSI and PSII, by determining the electron transport rate parameters ETR(I) and ETR(II) respectively (*33*), in mature wild-type and *sp2* mutant plants (Fig. 6F and G). Even though *sp2* plants do not display obvious visible differences from wild type under the normal conditions used, the mutant showed clearly elevated ETR(II) values, implying that CHLORAD normally acts to limit PSII activity.

**Fig. 6.**
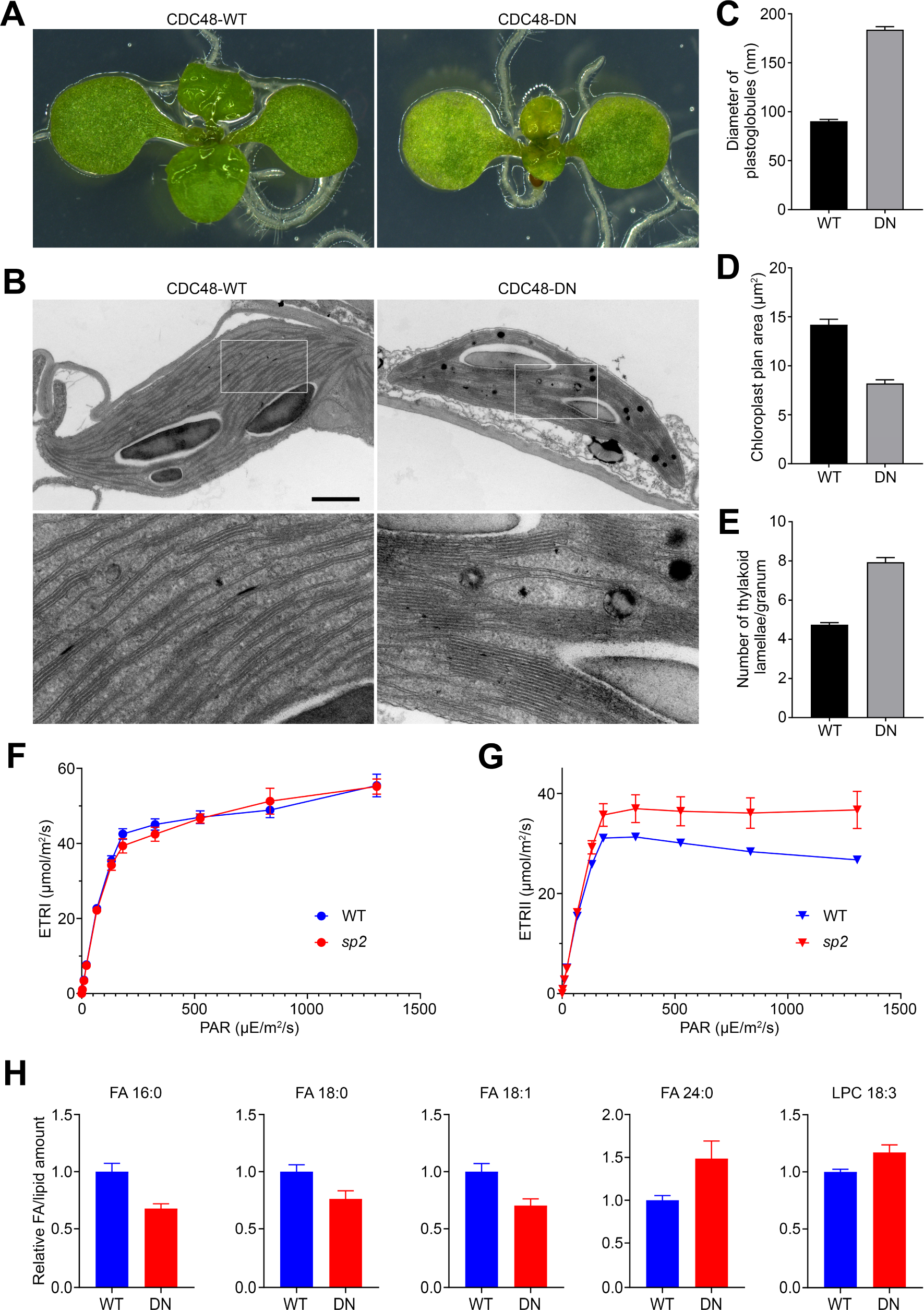
CHLORAD is required for normal photosynthetic function and lipid homeostasis. (**A** to **E**) Chloroplast development in CDC48-WT and CDC48-DN seedlings, after estradiol induction, was studied. Cotyledons of typical plants (A) were analysed by transmission electron microscopy (B), and representative images are shown. In B, the upper images are at the same magnification (scale bar, 2 μm); and below these are high magnification (4×) images corresponding to the boxed regions above. These and similar micrographs of chloroplasts were used to determine plastoglobule diameter (C), chloroplast cross-sectional area (D), and the number of thylakoid lamellae per granum (E) in each genotype. Values are means ± SEM from 50 chloroplasts. (**F** and **G**) Photosynthetic electron flow through PSI (ETRI) (F) and PSII (ETRII) (G) in wild-type and *sp2* mutant plants was measured. Plants were dark adapted before exposure to actinic light (PAR, photosynthetically active radiation) of the indicated intensities, followed by saturation pulses for the calculation of the indicated parameters. Values are means ± SEM from 5-6 plants. (**H**) Free fatty acid (FA) and polar lipid species in CDC48-WT and CDC48-DN seedlings, after estradiol induction, were quantified. Data were normalized to internal standards and per unit fresh weight; and the normalized values were expressed relative to CDC48-WT (taken as 1). The numbers alongside the FA/lipid names describe the constituent FA chains, in the format (number of carbons in the FA chain) : (number of double bonds in FA chain). Selected FA/lipid species showing significant differences (*P* < 0.05) in CDC48-DN relative to CDC48-WT are shown; a complete lipidomics dataset is provided in figure S6. Values are means ± SEM from 6 biological replicates. LPC, lysophosphatidylcholine.

Another critical role of chloroplasts is in lipid metabolism, as they are the major site of FA synthesis in plants (*34, 35*). Two chloroplast envelope proteins involved in FA metabolism, FAX1 and LACS9, were identified as candidate CHLORAD substrates via ubiquitinomics (table S3) or quantitative proteomics (table S5). Both proteins are involved in FA export to the ER, and changes in their expression influence FA and lipid homeostasis (*36, 37*), suggesting a role for CHLORAD in these processes. To investigate this possibility, we compared lipidomic profiles of CDC48-DN and CDC48-WT leaves, focusing on free FAs, glycerolipids, and glycerophospholipids. In total, 24 FA and lipid species were detected at significantly different levels in CDC48-DN (Student’s t test, *P* < 0.05). We observed decreases in chloroplast-produced FAs (16:0, 18:0, 18:1) and an increase in ER-elongated FA (24:0) (Fig. 6H, fig. S9, table S9), supporting the hypothesis that CHLORAD affects FA trafficking to the ER. Most lipid species were significantly reduced in CDC48-DN plants, with the exception of lysophosphatidylcholine (LPC) (18:3) and phosphatidylglycerol (PG) (16:0/18:1) (Fig. 6H, fig. S9, table S9), which may reflect a general disruption of FA and lipid homeostasis linked to the loss of normal regulation by CHLORAD.

## Discussion

Until recently, chloroplast proteolysis was thought to be dominated by internal chloroplast proteases of endosymbiotic origin (*3, 4*). Then, our discovery of CHLORAD revealed an important role for the UPS in the degradation of OEM proteins (*8, 9*). Internal chloroplast proteins were not considered to be likely targets of the cytosolic UPS, and of CHLORAD, due to the physical barrier presented by the double-membrane envelope. However, this work has now revealed how proteins of the chloroplast interior, including those in the IEM, stroma and thylakoids, can indeed be targeted by CHLORAD.

Proteomic evidence that chloroplast proteins are ubiquitinated has been presented previously (*28, 38, 39*). However, the relevant studies failed to determine whether the detected ubiquitination affects proteins that are resident in the organelle, which is highly pertinent given that most chloroplast proteins are synthesized in the cytosol before being imported into the organelle (*1, 11*). In fact, the identified ubiquitinated proteins were most likely cytosolic preproteins, for three reasons. First, the proteins were extracted from whole seedling or leaves, rather than from purified chloroplasts. Second, it is well-known that unimported chloroplast preproteins in the cytosol are processed by the UPS via an Hsp70/CHIP-dependent pathway (*27, 40*). And third, no chloroplast-encoded proteins were identified as being ubiquitinated in those studies, which is especially significant given that such proteins are extremely abundant.

Here, we conclusively demonstrated that chloroplast-resident proteins, including those of internal chloroplast compartments, are ubiquitinated and degraded by the proteasome. Evidence for internal ubiquitination included thermolysin treatment of purified chloroplasts and chloroplast subfractionation. Furthermore, ubiquitinomic analysis of isolated chloroplasts identified 316 chloroplast proteins, many from internal compartments; and of these 13 were chloroplast-encoded proteins, ruling out the possibility that they were modified in the cytosol as precursors. Those results were supported by quantitative proteomics, steady-state protein abundance and turnover data, and physiological analyses, which all together robustly demonstrated that ubiquitination affects the chloroplast’s proteome and functions extensively. Apart from the well-established targets in the TOC apparatus (*8, 9*), the substrates of CHLORAD identified here are involved in diverse processes including photosynthesis, fatty acid metabolism, and chloroplast gene expression.

An obvious question arising from these results concerns how internal chloroplast proteins are delivered to the cytosol for degradation. A retrotranslocation system comprising SP2 and CDC48 was previously shown to mediate the extraction of OEM substrates (*9*), and we present evidence here that the same components act on internal substrates too. First, the steady-state levels and turnover rates of internal chloroplast proteins were affected by SP2 and CDC48. Second, internal chloroplast proteins interacted physically with SP2 and CDC48. Third, in vivo retrotranslocation assays showed that CDC48 is required (presumably as a molecular motor) for the extraction of ubiquitinated internal proteins to the cytosol. However, exactly how stromal and thylakoidal proteins traverse the IEM remains unknown, as both SP2 and CDC48 operate at the OEM. This issue highlights a fundamental difference between ERAD (*15, 16*), which acts at a single membrane, and CHLORAD, which acts on endosymbiotic organelles with two bounding membranes. Although inner membrane retrotranslocation systems have not been identified, in either chloroplasts or mitochondria, our results indicate that such a system must exist in chloroplasts, and that there may even be internal ubiquitination mechanisms.

Photosynthesis, and its regulation, has been studied intensively for decades owing to its fundamental importance and iconic status as a defining feature of plants. Proteolysis is one of the key mechanisms for maintaining photosynthetic performance (*3, 4*). Turnover of internal chloroplast proteins involves multiple protease types, including three of prokaryotic origin: FtsH, Deg and Clp. For example, PSII core subunit D1 is frequently degraded by FtsH and Deg, as its reaction centre role renders it highly susceptible to photooxidative damage, necessitating frequent replacement. Other PSII core subunits, such as PsbB, PsbC and D2 (PsbD), are more stable and might only be degraded under stress conditions, although the mechanisms involved are less clear (*41, 42*). Even less information exists concerning the degradation of PSI subunits. The present study sheds light on PSI and PSII degradation, as it reveals an important role in photosynthetic regulation for the most pervasive proteolytic system in eukaryotes, the UPS, which is mediated by targeting key photosystem proteins including PsaA and PsbC.

Among the 24 photosystem (PSI and PSII) components identified in our ubiquitinome analysis, six are synthesized by the chloroplast itself. Analyses of the ubiquitination and turnover of PsaA and PsbC identified these proteins as bona fide CHLORAD substrates, while the importance of such regulation was supported by data on photosynthetic performance and chloroplast ultrastructure in CHLORAD mutants. Why photosystem core components are degraded by both organellar proteases and the UPS is unclear. One possibility is that the proteases function mainly in damage-induced responses whereas the UPS acts on specific substrates. Alternatively, the different systems might act at different stages or in response to different cues. For example, D1 requires frequent turnover in the light, whenever photosynthesis is active (*5-7*), and so thylakoid-localized proteases may be more economical and responsive to nearby chloroplast signals. In contrast, CHLORAD may be more suitable for the regulation of stable substrates that require infrequent degradation under conditions communicated by extrachloroplastic signals.

Circumstances warranting CHLORAD action may include abiotic stress (*14*) or those developmental phases during which internal organelle membranes are remodelled, such as de-etiolation (when the light-triggered degradation of prolamellar bodies occurs in etioplasts) or fruit ripening (when the disassembly of thylakoids occurs in developing chromoplasts) (*8, 17*). It is evident that protein degradation is crucial in these situations, but the underlying mechanisms are unclear. Our previous work revealed that CHLORAD is involved, by regulating the protein import machinery (*12, 13*); however, the new results reported here indicate that CHLORAD can also act much more directly, and that its established role in import regulation was arguably just the tip of the iceberg. For example, the maintenance of photosynthetic performance under senescence-inducing conditions in *sp1* and *sp2* mutants could be a consequence of reduced degradation of the photosystems, in addition to altered protein import (*8, 9*). Aside from the photosystem components, a number of other chloroplast-encoded photosynthesis proteins (i.e., one cytochrome b_6_f subunit, three ATP synthase subunits, one NDH complex subunit, and the Rubisco large subunit) were found to be ubiquitinated. Thus, it appears that CHLORAD can regulate diverse aspects of photosynthesis, including electron transport, energy transduction, and carbon fixation.

Ubiquitination sites were identified in 21 chloroplast-localized proteins with FA/lipid-related functions, while the abundance of six such proteins was affected by CDC48. These data point to an important and direct role for CHLORAD in the regulation of plant FA/lipid metabolism. One of the relevant proteins is FAX1, an IEM protein involved in FA export from chloroplasts that is required for normal cellular FA and lipid homeostasis. In *fax1* mutants, chloroplast-synthesized FAs (C16-18) are increased whereas very-long-chain FAs (which are elongated in the ER and thus require the export of C16-18 FAs) are reduced, with various developmental consequences (*37*). Our results identify FAX1 as a target of CHLORAD, and link the impairment of CHLORAD-dependent FAX1 degradation to perturbations in cellular FA/lipid homeostasis. Moreover, the pattern FA disturbances observed in CDC48-DN plants was strikingly similar to that seen previously in FAX1-overexpressing plants (*37*). It is important to note that carbohydrate metabolism and lipid metabolism are closely connected, and that the flow of carbon into oil (via FAX1) must be strictly controlled (*43*). Our results point to an important role for CHLORAD at this crucial metabolic nexus, the regulation of which is not currently well understood.

In this work we systematically identified the targets and ubiquitination sites of CHLORAD. Our results reveal that CHLORAD acts directly on a diversity of chloroplast cargos, extending far beyond the TOC apparatus to the organelle’s interior, and that the UPS broadly influences chloroplast functions including photosynthesis and lipid metabolism. Furthermore, the data illustrate how chloroplast proteostasis is remarkably complex and chimeric in nature, incorporating ancient systems inherited from the ancestral endosymbiont as well as eukaryotic, ubiquitin-dependent processes. These findings also provide new opportunities for cultivated plant improvement (e.g., by maximizing photosynthetic activity) (*44*), and thus may contribute to global challenges such as food security and carbon neutrality.

## Materials and Methods

### Plant growth conditions

All *Arabidopsis thaliana* plants were of the Columbia-0 (Col-0) ecotype. The *sp2-4* mutant, and transgenic lines expressing dominant-negative mutant and wild-type control versions of CDC48 with a C-terminal FLAG tag driven by an estradiol-inducible promoter (CDC48-DN and CDC48-WT, respectively), and SP2 with a C-terminal 6×Myc tag driven by the 35S promoter (SP2-6Myc), have all been described previously (*9*). All plants were grown under a long-day cycle (16 h light, 8 h dark), essentially as described previously (*45*). For in vitro growth, seeds were surface sterilized, sown on Murashige-Skoog (MS) agar medium in petri plates, cold-treated at 4°C, and thereafter kept in a growth chamber, as described previously (*45*). For the induction of CDC48-DN or CDC48-WT expression in the corresponding transgenic lines, 8-day-old plants were transferred onto MS medium supplemented with 4 μM estradiol (Sigma) and incubated for an additional two days.

### Measurement of photosynthetic electron flow

Photosynthetic parameters ETR(I) and ETR(II), which indicate electron flow through PSI and PSII, were measured on developmentally equivalent mature leaves (the third pair of true leaves) of 5-week-old plants using a DUAL-PAM-100 (Walz) in the Fluorescence and P700 Measure Mode of a dual channel. For the measurements, plants were dark adapted for 20 min at 22°C, before the actinic light intensity was increased stepwise from 0 to 1,300 μmol photons m^−2^ s^−1^. After exposure to each light intensity for 1 min, a 0.8 s pulse of saturating light (8,000 μmol photons m^−2^ s^−1^) was applied and the maximum PSII and PSI quantum yields, *F*_m_’ and *P*_m_’, respectively, were simultaneously recorded. Then, the stable PSII and PSI quantum yields, *F*_s_ and *P*_s_, were concurrently recorded. ETR(I) and ETR(II) were calculated as: 0.42 × PFD × (*P*_m_’ - *P*_s_) / *P*_m_ and 0.42 × PFD × (*F*_m_’ - *F*_s_) / *F*_m_’, respectively, where PFD is the photon flux density of actinic light, as described previously (*46*). Three experiments were performed, and approximately five leaves (each one from a different plant) were analysed per genotype in each experiment.

### Plasmid constructs

All primers used are listed in table S10. The *SP1-HA*, *YFP-HA*, *FLAG-Ub*, *CDC48-Trap-HA*, *Tic110-YFP* and *Toc33-HA* constructs have all been described previously (*8, 9*). All *Arabidopsis* CDSs were PCR-amplified from Col-0 cDNA. The Gateway cloning system (Invitrogen) was used to make most of the constructs, and all entry clones were verified by DNA sequencing. Constructs for the estradiol-inducible expression of untagged CDC48-DN or CDC48-WT (*CDC48-DNa* and *CDC48-WTa*, respectively) were generated using the pMDC7 binary vector (*47*), essentially as described previously (*9*) except without addition of the FLAG tag. To generate N-terminally 6×Myc-tagged ubiquitin (6Myc-Ub), the ubiquitin CDS was amplified from the *AtUBQ11* gene (At4g05050) and cloned into the pE3n vector (*48*), and then subcloned into the pB2GW7 35S-driven expression vector (*49*) for stable plant transformation (generating the *6Myc-Ub* construct). The CDSs of LACS9 (At1g77590), FAX1 (At3g57280), and CP12 (At3g62410) were cloned into the p2GWY7 plant expression vector (*49*) providing a C-terminal YFP tag (generating the *LACS9-YFP*, *FAX1-YFP* and *CP12-YFP* constructs). To generate haemagglutinin (HA)-tagged PrfB3 (At3g57190), the corresponding CDS was cloned into a modified p2GW7 vector (*9*) providing a C-terminal HA tag (generating the *PrfB3-HA* construct).

### Transient assays and stable plant transformation

Protoplast isolation and transient assays were carried out as described previously (*9, 50*). When required, bortezomib (Selleckchem) (prepared as a 10 mM stock solution in DMSO) was added to the protoplast culture medium at 15 h following transfection, to a final concentration of 5 μM; subsequently, the culture was incubated for a further 2-3 h before analysis. When using protoplasts isolated from the CDC48-WT and CDC48-DN transgenic lines, 10 μM estradiol (prepared as a 10 mM stock solution in ethanol) was included in the culture medium throughout the incubation of protoplasts. For YFP fluorescence and immunoprecipitation assays, 0.1 mL (10^5^ cells) or 1 mL (10^6^ cells) aliquots of protoplasts were transfected with 5 μg or 100 μg of DNA, respectively; the fluorescence signals were analysed after 15-18 h.

Transgenic lines carrying the *CDC48-DNa*, *CDC48-WTa* and *6Myc-Ub* constructs were generated by *Agrobacterium*-mediated transformation (*51, 52*). Transformants were selected using MS medium containing 50 μg/mL hygromycin B (Melford) or 10 μg/mL phosphinotricin (Duchefa). Approximately ten independent T2 lines were analysed per construct, and one representative line with a single T-DNA insertion (which showed a 3:1 segregation on selective MS medium in the T2 generation) were chosen for further analysis.

### Microscopy

Transmission electron microscopy was performed as described previously (*51*). For induction of CDC48-WT or CDC48-DN expression in the corresponding transgenic lines, 8-day-old plants were transferred onto MS agar medium supplemented with 4 μM estradiol and incubated for an additional two days. Measurements were recorded using at least 50 different plastids per genotype, and were representative of three individuals per genotype. Chloroplast cross-sectional area was estimated as described previously (*51, 53*), using the equation: π × 0.25 × length × width. Numbers of thylakoid lamellae per granal stack were counted as previously described (*8, 51*) for at least 180 resolvable grana from three individuals per genotype. Plastoglobule diameter was determined by measuring at least 150 different plastoglobules from three individuals per genotype.

All fluorescence microscopy experiments were conducted at least twice with the same results, and typical images are presented. The imaging of YFP and chlorophyll fluorescence signals was conducted by examining protoplasts using a Zeiss LSM 510 META laser-scanning confocal microscope (Carl Zeiss Ltd.), as described previously (*14*). All images were captured using the same settings to enable comparisons.

### Chloroplast isolation, protease treatment, and subfractionation

Chloroplasts were isolated from plants grown in vitro for 8 to 10 days (or, when stated, from protoplasts). For CDC48-DN and CDC48-WT, 8-day-old plants were transferred onto MS agar medium (without sucrose) supplemented with 4 μM estradiol and incubated for an additional two days before chloroplast isolation. For 6Myc-Ub, 8-day-old plants were transferred to MS liquid medium (without sucrose) supplemented with 5 μM bortezomib and incubated for an additional two days before chloroplast isolation. Chloroplast isolations and protease treatments were performed as described previously (*9, 45*).

For chloroplast subfractionation, pelleted chloroplasts from 6Myc-Ub plants were first resuspended and lysed in prechilled hypotonic lysis buffer (25 mM HEPES-KOH, pH 8.0, supplemented with 0.5% plant protease inhibitor cocktail [PPIC, Sigma, P9599]) in a rotator for 30 min at 4°C. Then, the lysate was centrifuged at 18,000 × *g* for 30 min at 4°C. The recovered supernatant was centrifuged again at 18,000 × *g*, for 30 min at 4°C, and the resulting supernatant (S18) was retained for further analysis. The pellet of the first centrifugation step was resuspended in prechilled hypotonic lysis buffer, and centrifuged at 18,000 × *g* for 30 min at 4°C. The pellet resulting from this centrifugation step (P18) was retained for further analysis.

### SDS-PAGE, immunoblotting and immunoprecipitation

Sodium dodecyl sulphate-polyacrylamide gel electrophoresis (SDS-PAGE) and immunoblotting were performed essentially as described before (*53, 54*). Primary antibodies were as follows. To detect TOC proteins or components of the translocon at the inner envelope membrane of chloroplasts (TIC), we employed: anti-atToc75-III antibody (*52*); anti-atToc159 antibody (*55*); anti-atToc33 (G-domain) antibody (*52*); anti-atTic110 antibody (*56, 57*); and anti-atTic40 antibody (*52*). To detect non-TOC outer envelope membrane proteins, we employed: anti-CHUP1 antibody (*58*); and anti-FtsZ2 antibody (*59*). To detect chloroplast stromal proteins, we employed: anti-RPL2 (HUABIO, PAB20010; or as described previously (*60*)); anti-COR15 antibody (*61*); anti-RbcL antibody (*62*); and anti-PAO antibody (*63*). To detect chloroplast thylakoid proteins, we employed: anti-PsaA antibody (HUABIO, PAB01001; or as described previously (*64*)); anti-PsbC (CP43) antibody (HUABIO, PAB02003); anti-PsbD antibody (Agrisera AS06146); and anti-LHCP antibody (*51, 52*). To detect proteins of other cellular compartments, we employed: anti-α-tubulin (cytosol; HUABIO, T5168); and anti-H3 histone (nucleus; Abcam, ab1791). Other primary antibodies we employed were: anti-HA tag antibody (Sigma, H6908); anti-c-Myc tag antibody (Abcam, ab9106); anti-GFP antibody (detects both GFP and YFP; Sigma, SAB4301138); and anti-FLAG tag antibody (Sigma, F7425). We used Tic110, α-tubulin or H3 as loading controls.

Secondary antibodies were polyclonal goat anti-rabbit IgG conjugated with horseradish peroxidase (Sigma, 12-348), or, in the case of immunoprecipitations, monoclonal mouse anti-rabbit IgG light-chain-specific conjugated with peroxidase (Jackson ImmunoResearch, 211-032-171). Chemiluminescence was detected using EZ-ECL (Biological Industries, Sartorius) or ECL Plus Western Blotting Detection Reagents (GE Healthcare), and an LAS-4000 imager (Fujifilm). Band intensities were quantified using ImageJ (*65*) or Aida software (Raytest). Quantification data were based on results from at least three experiments all showing a similar trend. Typical images are shown in all figures.

For the immunoprecipitation of HA-tagged proteins, total protein (∼500 mg) was extracted from protoplasts in IP buffer (25 mM Tris-HCl, pH 7.5, 150 mM NaCl, 1 mM EDTA, 1% Triton X-100) containing 0.5% PPIC (Sigma), and centrifuged at 20,000 × *g* for 10 min at 4°C. The clear lysate was then incubated with 50 μL EZview Red Anti-HA Affinity Gel (Sigma) for 2 h to overnight at 4°C with slow rotation. After six washes with 500 μL IP-washing buffer (25 mM Tris-HCl, pH 7.5, 150 mM NaCl, 1 mM ethylenediaminetetraacetic acid [EDTA], 0.5% Triton X-100), bound proteins were eluted by boiling in 2× SDS-PAGE loading buffer (50 mM Tris-HCl, pH 6.8, 20% glycerol, 1% sodium dodecyl sulphate [SDS], 0.1 M dithiothreitol [DTT]) for 5 min, and analysed by SDS-PAGE and immunoblotting. A similar procedure was adopted for the immunoprecipitation of Myc- or FLAG-tagged proteins, except that 50 μL EZview Red Anti-c-Myc Affinity Gel (Sigma) or Anti-FLAG M2 Affinity Gel (Sigma) was used instead of the anti-HA gel. When detecting ubiquitinated proteins, the IP buffer also contained 10 mM N-ethylmaleimide (NEM; Sigma).

To assess in vivo ubiquitination of chloroplast substrates, FLAG-Ub was transiently overexpressed in protoplasts to increase detection sensitivity for higher molecular weight forms (*9*). Protoplasts were lysed in denaturing buffer (25 mM Tris-HCl, pH 7.5, 150 mM NaCl, 5 mM EDTA, 10 mM NEM, 1% SDS, 2% Sarcosyl, 5 mM DTT), prior to incubation at 75°C and 600 rpm for 30 min in a Thermomixer Comfort (Eppendorf). The lysate was then diluted by adding 1 volume of 2% Triton X-100 and 8 volumes of IP buffer containing 0.5% PPIC, and incubated on ice for 30 min. Ubiquitinated proteins were enriched through subsequent immunoprecipitation steps as described above with Anti-FLAG M2 Affinity Gel. Immunoblot analysis of the precipitates using antibodies against proteins of interest was used to demonstrate protein ubiquitination, as indicated by higher molecular weight smears.

### Mass spectrometry analysis of Myc-tagged ubiquitin interactors

Eight-day-old 6Myc-Ub plants were transferred to liquid MS medium (without sucrose) supplemented with 5 μM bortezomib, and incubated for an additional two days before chloroplast isolation. Isolated chloroplasts were subjected to immunoprecipitation using EZview Red Anti-c-Myc Affinity Gel (see above). Elution of immunoprecipitated proteins was performed using 100 μg/mL c-Myc peptide (Sigma, M2435), on ice for 15 minutes, and eluates were collected following centrifugation at 8,200 × *g* for 30 s at 4°C.

Protein digestion was performed using a filter-aided sample preparation protocol. Eluate (50 μL) was denatured and alkylated with 950 μL urea buffer (8 M urea, 100 mM ammonium bicarbonate [AB], 10 mM Tris[2-carboxyethyl]phosphine hydrochloride, 50 mM 2-chloroacetamide, 0.2% protease inhibitor [Sigma, P8340]), and incubated in the dark at room temperature for 30 min. Filters (30 kD cut-off, Vivacon 500) were pre-washed (using 0.1% [v/v] trifluoroacetic acid [TFA], 50% [v/v] acetonitrile), and then loaded with 200 μL of 8 M urea in 100 mM AB, followed by 100 μL of the denatured sample and an additional 100 μL of 8 M urea in 100 mM AB. Filters were centrifuged at 14,000 × *g* for 15 min, and then washed twice using 6 M urea in 25 mM AB. LysC protease (Promega, V1671; 200 ng in 50 μL of 6 M urea, 25 mM AB) was added to each filter, prior to incubation at 37°C for 4 h. The digestion solution was diluted to <1 M urea upon addition of trypsin (Promega, V511A; 200 ng in 300 μL of 1 M urea, 25 mM AB, 1 mM CaCl2), prior to incubation at 37°C overnight. After digestion, the eluate fractions were collected by centrifugation at 14,000 × *g* for 15 min at 25°C. The filters were washed using 0.1% TFA, and then 0.1% TFA, 50% acetonitrile, and the flow-through fractions were combined with the initial eluate fractions and dried down in a SpeedVac concentrator.

Resulting tryptic peptides were analysed on an EASY-nLC 1000 system (Thermo Fisher) connected to a Q Exactive mass spectrometer (Thermo Fisher) through an EASY-Spray nano-electrospray ion source (Thermo Fisher). The peptides were initially trapped on a C18 PepMap100 pre-column (300 µm i.d. x 5 mm, 100 Å, Thermo Fisher) using solvent A (0.1% formic acid in water). Trapped peptides were separated on a in-house constructed analytical column (75 µm i.d. × 500 mm, Reprosil C18, 1.9 µm, 100 Å) using a linear gradient (length: 60 minutes, 18 % to 30 % solvent B [0.1% formic acid, 5 % DMSO in acetonitrile]; flow rate: 200 nL/min). The separated peptides were electrosprayed directly into the mass spectrometer operating in a data-dependent mode. Full scan MS spectra were acquired in the Orbitrap (scan range 350-1500 m/z, resolution 70000, AGC target 3e6, maximum injection time 50 ms). After the MS scans, the ten most intense peaks were selected for HCD fragmentation at 30% of normalised collision energy. HCD spectra were also acquired in the Orbitrap (resolution 17500, AGC target 5e4, maximum injection time 120 ms).

Protein identification and quantification were performed using the Andromeda search engine implemented in MaxQuant (1.6.3.4). Peptides were searched against a reference proteome of *Arabidopsis thaliana* (Uniprot database, downloaded Jan 2017) with the custom addition of the 6×Myc-tagged ubiquitin sequence. The di-Gly-modification at lysine was added as an additional variable modification, while other settings were kept at default parameters (*66*). False discovery rate (FDR) was set at 1% for both peptide and protein identification.

### Di-Gly ubiquitinome analyses

Ubiquitinome analyses were performed using isolated chloroplast samples from wild-type and CDC48-DN plants. Wild type samples were analysed in the Advanced Proteomics Facility, University of Oxford, and CDC48-DN samples were analysed by Shanghai Applied Protein Technology Co., Ltd.

Isolated wild-type chloroplasts were lysed in urea buffer (8 M urea, 100 mM ammonium bicarbonate, 10 mM Tris[2-carboxyethyl]phosphine hydrochloride, 50 mM 2-chloroacetamide) containing 1% PPIC at room temperature for 30 min in dark. The lysate was cleared by centrifugation at 4,500 × *g* for 15 min at room temperature, and then the concentration of urea in the supernatant was diluted to 6 M using 25 mM ammonium bicarbonate. LysC protease (Wako, 129-02541) was added (to a LysC:protein ratio of 1:100 [w/w]), and digestion was performed at 37°C for 4 h with mixing. The urea concentration was diluted to 1 M using 25 mM ammonium bicarbonate, then CaCl_2_ was added to a final concentration of 1 mM. Next, Trypsin (Sigma, T-1426) was added (to a trypsin:protein ratio of 1:40 [w/w]), and digestion was performed at 37°C overnight with mixing. Trifluoroacetic acid was added (to a final concentration of 1%), precipitation was allowed to occur for 15 min on ice, and the precipitates were removed by centrifugation at 1,780 × *g* for 15 min at room temperature. Peptides in the supernatant were then purified using Sep-Pak C18 columns (Waters, WAT051910). For analysis of di-Gly modification-enriched peptides, an immunoprecipitation step was performed using a PTMScan Ubiquitin Remnant Motif (K-ε-GG) Kit (Cell Signal Technology) according to the manufacturer’s instructions.

LC-MS/MS analysis was conducted as described above (for the analysis of Myc-tagged ubiquitin interactors), with the following changes: The peptides were analysed on a nanoUHPLC (Thermo Fisher). Trapped peptides were separated on an EASY-Spray Acclaim PepMap analytical column (75 µm i.d. × 500 mm, RSLC C18, 2 µm, 100 Å; Thermo Scientific) using a linear gradient (length: 60 min, 15 % to 35 % solvent B [0.1% formic acid, 5 % DMSO in acetonitrile]; flow rate: 200 nL/min).

Protein identification and quantification were performed using Sequest HT in Proteome Discoverer 1.4 (Thermo Fischer, version 1.4.0.288). Tandem mass spectra were searched against a database containing 16480 protein entries from *Arabidopsis thaliana* (Uniprot, release from July 2022) and common contaminants. During database searches, it was considered that: cysteines were fully carbamidomethylated (+57.0215, statically added), methionines were fully oxidized (+15.9949, dynamically added), all N-terminal residues were acetylated (+42,0106, dynamically added), and lysines were ubiquitinated (di-Gly +114.043, dynamically added). Two missed cleavages were permitted. FDR was set at 1% for both peptide and protein identification. Protein identification and quantification were also conducted as described above (for the analysis of Myc-tagged ubiquitin interactors), without the custom addition of the 6×Myc-tagged ubiquitin sequence to the database.

For analysis of di-Gly modification-enriched peptides of CDC48-DN samples, isolated chloroplasts were lysed in urea buffer (8 M urea, 100 mM Tris-HCl, pH 8.5) for protein extraction. Then, DTT was added to a final concentration of 10 mM, and the samples were mixed at 600 rpm for 1.5 h at 37°C, and cooled to room temperature before being subjected to trypsin digestion and desalting. Desalted peptides (∼ 2 mg) were reconstituted in 1.4 mL precooled IAP buffer (50 mM MOPS, pH 7.2, 10 mM sodium phosphate, 50 mM NaCl), and then incubated with anti-K-ε-GG antibody beads (PTMScan Ubiquitin Remnant Motif [K-ε-GG] Kit, Cell Signal Technology) at 4°C for 1.5 h. The antibody beads were collected by centrifugation at 2,000 × *g* for 30 s, and then washed three times with 1 mL precooled IAP buffer and three times with precooled water. The di-Gly peptides were eluted with 40 μL 0.15% TFA after incubation at room temperature for 10 min, the elution was repeated once, and the eluates were combined. After centrifugation at 2,000 × *g* for 30 s, supernatants were collected and desalted using Millipore C18 StageTips (Sigma, ZTC18S096).

LC-MS/MS analysis of CDC48-DN samples was performed using a Q Exactive mass spectrometer (Thermo Fisher) that was coupled to an EASY-nLC (Thermo Fisher) for 60, 120 or 240 min. The peptides were loaded onto a reversed phase trap column (Acclaim PepMap100, 100 μm × 2 cm, nanoViper C18; Thermo Scientific) connected to the C18-reversed phase analytical column (EASY-Column, 10 cm long, 75 μm inner diameter, 3 μm particle size; Thermo Scientific) in buffer A (0.1% formic acid) and separated with a linear gradient of buffer B (84% acetonitrile and 0.1% formic acid) at a flow rate of 300 nL/min controlled by IntelliFlow technology. The mass spectrometer was operated in positive ion mode. MS data were acquired using a data-dependent top10 method, dynamically choosing the most abundant precursor ions from the survey scan (300-1800 m/z) for high-energy collisional dissociation (HCD) fragmentation. Automatic gain control (AGC) target was set to 3 × 10^6^, and maximum inject time was set to 10 ms. Dynamic exclusion duration was 40.0 s. Survey scans were acquired at a resolution of 70,000 at m/z 200, resolution for HCD spectra was set to 17,500 at m/z 200, and isolation width was 2 m/z. Normalized collision energy was 30 eV and the underfill ratio, which specifies the minimum percentage of the target value likely to be reached at maximum fill time, was defined as 0.1%. The instrument was run with peptide recognition mode enabled.

The MS raw data for each sample were combined and searched using the MaxQuant software (*67*) against an *Arabidopsis* UniProt database (*68*) for chloroplast (plastid)-associated proteins. Trypsin was chosen as the enzyme with a maximum of two missed cleavages allowed. Precursor and fragment mass error tolerances were set at 20 ppm and 0.1 Da, respectively. Peptide modifications allowed during the search were: carbamidomethyl (C, fixed), GlyGly (K, variable), and oxidation (M, variable). All matched MS/MS spectra were filtered by mass accuracy and matching scores to reduce protein FDR to ≤1%, based on the target-decoy strategy using a reversed database.

### Quantitative proteomic analysis

Eight-day-old CDC48-DN and CDC48-WT plants were transferred onto MS agar medium supplemented with 4 μM estradiol and incubated for a further two days before whole seedlings were collected, weighed and frozen in liquid N_2_. Three independent samples of each genotype were collected. Quantitative proteomic analysis was performed by Shanghai Luming Biotechnology Co., Ltd. Frozen seedling samples (500 mg each) were ground to a power and then further ground in 1 mL extraction buffer (250 mM Tris-HCl, 0.7 M sucrose, 100 mM NaCl, 50 mM EDTA, 10 mM DTT, pH 7.8; freshly prepared). The samples were then mixed with an equal volume of Tris-buffered phenol for 30 min at 4°C, before centrifugation at 7,100 × *g* for 10 min at 4°C to collect the upper phenolic phase. The supernatants were mixed with five volumes of 0.1 M cold ammonium acetate / methanol solution, and precipitated at -20°C overnight. Then, the proteins were pelleted by centrifugation at 12,000 × *g* at 4°C for 10 min (all centrifugation steps below were similar). Pellets were washed twice with ice-cold methanol, and then twice more with ice-cold acetone; after each washing step, the samples were pelleted by centrifugation. The final pellets were dried at room temperature for 3 min and resuspended in SDS lysis buffer (Beyotime, Shanghai) for 3 h. Lastly, the samples were centrifuged two further times, and the supernatants were retained.

The filter-aided sample preparation method (*69*) was used for proteolysis. Briefly, 100 μg of each sample was placed in an Amicon ultrafiltration centrifuge tube (Merck, 10 kD molecular weight cut-off), 120 μL reductant buffer (10 mM DTT, 8 M urea, 100 mM triethylammonium bicarbonate [TEAB], pH 8.0) was added, and the mixture was incubated for 1 h at 60°C. Iodoacetamide was added to a final concentration of 50 mM, and the solution was incubated in darkness at room temperature for 40 min. After centrifugation into a collection tube, the flow-through solution was discarded. Then, 100 μL of 300 mM TEAB buffer was added to the ultrafiltration tube, followed by 3 μL sequencing-grade trypsin solution (1 μg/μL, Promega), and the sample was incubated for 12 h at 37°C. Digested peptides were collected as the flow-through in a new collection tube by centrifugation. Then, 50 μL of 200 mM TEAB solution was added to the ultrafiltration cartridge, and the flow-through solution was combined with the sample in the same collection tube after centrifugation. Digested peptides were desalted using a SOLA SPE 96-well Column (Thermo Fisher).

The LC-MS/MS analyses were performed on an Xcalibur 2.2 SP1 system coupled with a Q Exactive mass spectrometer (both Thermo Fisher). Peptide concentrations of all samples were adjusted to 20.6 ng/μL with 0.1% TFA. For each analysis, 10 μL of sample (corresponding to a total sample amount of 300 ng of proteins digested with trypsin) was injected. After injection, peptides were pre-concentrated with 0.1% TFA on a trap column (AcclaimR PepMap 100 column, 100 μm i.d. × 2 cm, C18, 5 μm particle size, 100 Å pore size; Thermo Scientific) at a flow rate of 7 μL/min for 10 min. Subsequently, the analyte was transferred to the analytical column (AcclaimR PepMap RSLC column, 50 μm i.d. × 15 cm, C18, 2 μm particle size, 100 Å pore size; Thermo Scientific) and separated at 60°C using a 90 min gradient from 5% to 95% solvent B at a flow rate of 220 nL/min (solvent A: 0.1% formic acid, solvent B: 0.1% formic acid in acetonitrile). The gradient elution conditions were as follows: 0-55 min, 5% B to 30% B; 55-80 min, 30% B to 50% B; 80-90 min, 50% B to 100% B. The mass spectrometer was operated in a data-dependent mode. Full-scan mass spectra were acquired at a mass resolution of 70,000 (mass range 350-2000 m/z) in the Orbitrap analyser. Tandem mass spectra of the 20 most abundant peaks were acquired in the linear ion trap by peptide fragmentation using collision-induced dissociation (CID). Normalized collision energy (NCE) was set to 27% and an isolation width of 2.0 m/z was chosen.

Analyses of MS/MS spectra followed a similar protocol to that described for Di-Gly ubiquitinome analysis, with the following differences: database searching was done using the *Arabidopsis* UniProt Reference Proteome with a 10 ppm mass tolerance for precursors. Relative protein quantification was performed using the MaxQuant software for label-free quantification analysis and the search engine was Andromeda. Quantitation was carried out only for proteins with two or more unique peptide matches.

### Annotation for subcellular and subplastidic localization and function

Proteins identified by LC-MS/MS were manually annotated with respect to subcellular and subplastidic localization, and function. Information on protein characteristics, function, and subcellular and subplastidic localization was searched for sequentially and manually in several public databases (TAIR (*70*), UniProt (*68*), NCBI (*71*)) and in the appropriate literature, in order to annotate well-characterized chloroplast proteins. Two specialist databases, PPDB (*22*) and AT_CHLORO (*23*), were used to assign chloroplast and subplastidic localizations for some less well-characterized proteins. For proteins not included in these databases, the ChloroP (*72*) and TargetP (*73*) prediction tools were used to test for the presence of a transit peptide (a typical feature of chloroplast proteins); or the SUBA4 database (*74*) was used, which provides information on subcellular localization from published proteomic or fluorescent protein fusion datasets. Proteins associated with the chloroplast envelope were assessed by reference to a recently published proteomics study (*18*). Chloroplast proteins with unclear subplastidic localization were analysed using the ARAMEMNON database (*75*) for the prediction of transmembrane domains, to rule out the possibility of stroma or intermembrane space localization. Functions of proteins were classified with gene ontology (GO) terms using the Protein Analysis THrough Evolutionary Relationship (PANTHER) Classification System (*76*).

### RNA sequencing

Eight-day-old CDC48-DN and CDC48-WT plants were transferred onto MS agar medium supplemented with 4 μM estradiol and incubated for a further two days before whole seedlings were collected, weighed and frozen in liquid N_2_. Three independent samples of each genotype were collected, and total RNA was extracted using TRIzol reagent (Invitrogen, Thermo Fisher). Sample RNA quality and integrity were assessed by agarose gel electrophoresis. RNA sequencing and data analysis were performed by Shanghai HanYu Biotech lab.

For RNA-seq library preparation and sequencing, RNA was purified using Dynabeads Oligo(dT)25 (Invitrogen, Thermo Fisher) followed by DNase I (Takara) treatment. Library preparation employed 100 ng RNA per sample, and was done using an NEBNext Ultra RNA Library Prep Kit for Illumina (NEB). After cluster generation on a cBot cluster generation system using Truseq PE Cluster kit (Illumina), the prepared libraries were sequenced on Illumina Nova, resulting in paired-end reads.

To obtain clean reads, raw data (raw reads) in FASTQ format were processed using Trimmomatic v0.32 to remove data containing poly-N and low-quality reads. The clean reads were then mapped to the *Arabidopsis* reference genome (*70*) using bowtie2 v2.1.0. The expression levels of genes were calculated in FPKM (Fragment Per Kilobase of gene per Million reads mapped). Differentially expressed genes were identified using the P package DEGseq v1.20.0. Genes with log2 fold change greater than 1 or less than -1, and adjusted *P* value lower than 0.001, were classified as differentially expressed genes.

### Protein degradation assays

Protein degradation was investigated by using cycloheximide-chase degradation assays or lincomycin-chase degradation assays, as described previously (*77, 78*) with modifications. Cycloheximide and lincomycin are translational inhibitors used to block the synthesis of nucleus-encoded proteins and chloroplast-encoded proteins, respectively.

To monitor degradation of PrfB3, wild-type protoplasts transfected with the *PrfB3-HA* construct were analysed by cycloheximide treatment. Transformed protoplasts were incubated for 15 h, and then cycloheximide was applied to a final concentration of 10 μg/mL (dissolved in water), before further incubation for the durations indicated in the figure legends. The UPS dependency of PrfB3 turnover was determined by applying cycloheximide with and without bortezomib, and samples were analysed after 6, 12 and 24 h of incubation.

For lincomycin treatment, seedlings grown in vitro for 8 days were transferred to MS liquid medium (without sucrose) containing 400 μM lincomycin (dissolved in water), and then further grown under normal conditions for the durations indicated in the figure legends. Seeding samples (50 mg fresh weight) were flash frozen in liquid N_2_ upon collection, and protein extracts were prepared as previously described (*53*). For the induction of the CDC48-DN and CDC48-WT constructs, 8-day-old transgenic plants were transferred to liquid MS medium containing 4 μM estradiol and incubated for 2 days; lincomycin was added during this incubation at 0, 12, 24 and 36 h to achieve the desired periods of treatment. Experiments were performed with three biological replicates, and for each replicate at least 10 seedlings were analysed per genotype at each time point.

### In vivo retrotranslocation assays

In vivo retrotranslocation assays were performed as previously described with minor changes (*9*). First, FLAG-Ub was transiently overexpressed in 10^6^ protoplasts of each genotype (CDC48-DNa and CDC48-WTa) to increase detection sensitivity for higher molecular weight (ubiquitinated) forms of the chloroplast substrates (*8*). The transformed protoplasts were incubated for 15 h, and then bortezomib was applied to a final concentration of 5 μM before an additional 3 h incubation. Subsequent fractionation steps to produce separate chloroplast and cytosol samples were all carried out on ice or at 4°C, and used previously described procedures with modifications (*9*). Protoplasts were pelleted by centrifugation at 100 × *g* for 2 min, and gently resuspended with protoplast-washing buffer (500 mM mannitol, 4 mM 4-morpholineethanesulfonic acid [MES]-KOH, pH 5.6). Then, the protoplasts were pelleted again and resuspended by gentle agitation in 500 μL HS buffer (50 mM 4-(2-hydroxyethyl)piperazine-1-ethanesulfonic acid [HEPES]-NaOH, pH 8.0, 0.3 M sorbitol) containing 0.5% PPIC and 5 μM bortezomib, and gently forced twice through 10 μm nylon mesh to release chloroplasts. The collected flow-through was centrifuged at 1,000 × *g* for 5 min to produce a chloroplast-containing pellet and a cytosol-containing supernatant (S1). The pellet was gently resuspended in 500 μL HS buffer, and the chloroplasts were purified using a two-step Percoll (Fisher Scientific) gradient (*45*). Intact chloroplasts were washed with 500 μL HS buffer, and then pelleted by centrifugation at 1,000 × *g* for 5 min. The S1 sample was centrifuged at 10,000 × *g* for 15 min. The resulting supernatant (S10) was recovered and ultracentrifuged at 100,000 × *g* for 1 h, producing a further supernatant (S100) that was concentrated to 50 μL by using Vivaspin 500 ultrafiltration spin columns; this was the cytosolic fraction. Subsequence detection of substrate ubiquitination in both chloroplast and cytosolic fractions was performed by immunoprecipitation (anti-FLAG) and immunoblotting as described above. Experiments were repeated three times, and similar results were obtained.

### Measurement of lipids and free fatty acids

Plant material was harvested as described above for quantitative proteomic analysis, except that for each genotype six independent samples were processed. Measurement of lipids and FAs was performed by Shanghai Luming Biotechnology Co., Ltd. Lipids/FAs were extracted as described by Folch et al. (*79*) with modifications. In brief, lipids/FAs were extracted from ∼50 mg frozen tissue powder using 1 mL chloroform:methanol (2:1, v/v), spiked with known amounts of isotope-labelled internal standards. The samples were vortexed for 30 s and then incubated at 4°C in an ultrasonic bath for 10 min. Subsequently, samples were placed at -20°C for 30 min, and in each case the lower organic phase was recovered and centrifuged at 13,000 × *g* for 10 min at 4°C. The supernatants were collected and the lipid extraction step was repeated. The cleared organic phases were combined and evaporated using a SpeedVac system. Finally, samples were reconstituted in isopropanol:methanol (1:1, v/v).

Liquid chromatography was performed as described previously (*80*), with modifications, using an ExionLC System (Sciex) consisting of a binary high pressure mixing gradient pump with degasser, a thermostated autosampler, and a column oven; the mass spectrometer was a QTRAP® 6500+ (SCIEX) equipped with an IonDrive Turbo V source. The temperature of the autosampler was set at 10°C. The mobile phases were acetonitrile:water (6:4, v/v) with 0.1% formic acid and 10 mM ammonium formate (eluent A), and acetonitrile:methanol (1:9, v/v) with 0.1% formic acid and 10 mM ammonium formate (eluent B). For each injection, a 5 μL sample was loaded into an Aquity UPLC HSS T3 column (2.1 mm i.d. × 10 cm, 1.7 µm particle size; Waters) using: a 1.5 min initial flow of 0% B; a 3.5 min linear gradient from 0% B to 55% B; a 5 min linear gradient from 55% B to 60% B; a 3 min linear gradient from 60% B to 70% B; a 2 min linear gradient from 70% B to 90% B; and a 1 min linear gradient from 90% B to 100% B, and held for 2 min. The flow rate was set at 350 μL/min. Finally, the buffer was set back to 0% B, and the column was re-equilibrated for 2 min. Thus, the total running time was 20 min.

The MS method was performed in positive and negative electrospray ionization (ESI) modes, working in the time-scheduled multiple reaction monitoring (MRM) method. The source conditions were as follows: curtain gas was 35 psi, collision gas was medium, the ion spray voltage was -4500 V/+5500 V, and ion source gas 1 and ion source gas 2 were 40 psi and 45 psi, respectively. To evaluate the stability of the system during the whole analysis process, quality control samples obtained by mixing equal volumes (20 μL) of each sample were inserted into the analysis queue and determined three times in both positive and negative ESI modes. Spectra data were collected using Analyst Software 1.7 (Sciex) according to the manufacturer’s instructions. Analysis of the spectra (alignment, peak picking, normalization, and peak integration) was performed using MRMPROBS (*81*). Lipids/FAs were identified by spectral and retention time matching with authentic compounds using an in-house custom library augmented with a library from Riken. Compounds were normalized relative to the internal standard.

### Statistical analysis

Statistical calculations (mean, standard error of the mean, t-test) were performed using GraphPad Prism software. Statistical significance of differences between two experimental groups was assessed by using a two-tailed Student’s t-test. Differences between two datasets were considered significant at *P* < 0.05.

## Acknowledgements

Dedicated to the memory of Danny J. Schnell. We thank X. Gao, Z. Zhang and J. Li for electron microscopy; M. Wada (CHUP1), K. W. Osteryoung (FtsZ2), J. Gray (PAO), K. Redding (PsaA), K. Yamaguchi (RPL2), R. Spreitzer (RbcL), N. E. Hoffman (LHCP), and T. Inaba (COR15) for antibodies; H. Zhang for the FAX1-YFP construct and technical assistance with confocal microscopy.

## Author contributions

RPJ, QL and YS conceptualized and designed the experiments. YS and ZY performed most of the experiments and contributed equally as co-first authors. QL set up the initial screen for CHLORAD substrates. HC performed IP-MS, supervised by SM. YY contributed to the IP experiments. YL performed protein degradation assays, supervised by YH and ZY. WB generated the 6Myc-Ub lines. GC performed the photosynthetic analysis. MF performed ubiquitinome assays on wild-type plants. RPJ and QL co-supervised the project and wrote the manuscript with input from all co-authors.

## Data and materials availability

The mass spectrometry proteomics data have been deposited to the ProteomeXchange Consortium via the PRoteomics IDEntifications (PRIDE) repository (https://www.ebi.ac.uk/pride/archive/) with the identifier pending, and to the Integrated Proteome Resources (iProX, https://www.iprox.cn/) with the identifier IPX0004051000. The RNA-seq data have been deposited to NCBI Sequence Read Archive (https://submit.ncbi.nlm.nih.gov/) with the identifier PRJNA801890. All other data are available in the manuscript or the supplementary materials.

## Supplementary Materials

### Tables

**Table S1.** Identification of chloroplast proteins that co-immunoprecipitate with 6Myc-Ub.

**Table S2.** Identification of ubiquitinated proteins in wild-type chloroplasts using di-Gly ubiquitinomics.

**Table S3.** Identification of ubiquitinated chloroplast proteins in CDC48-DN plants using di-Gly ubiquitinomics.

**Table S4.** Chloroplast-encoded proteins with ubiquitination sites.

**Table S5.** Quantitative proteomic analysis of chloroplast proteins in CDC48-DN and CDC48-WT plants.

**Table S6.** Selected putative CHLORAD substrates identified by quantitative proteomics.

**Table S7.** Quantitative proteomic analysis of isolated CDC48-DN and CDC48-WT chloroplasts.

**Table S8.** Quantitative transcriptomic analysis of CDC48-DN and CDC48-WT plants.

**Table S9.** Lipidomic analysis of CDC48-DN and CDC48-WT plants.

**Table S10.** Primers used during the course of the study.

### Supplementary Figures

**Fig. S1.**
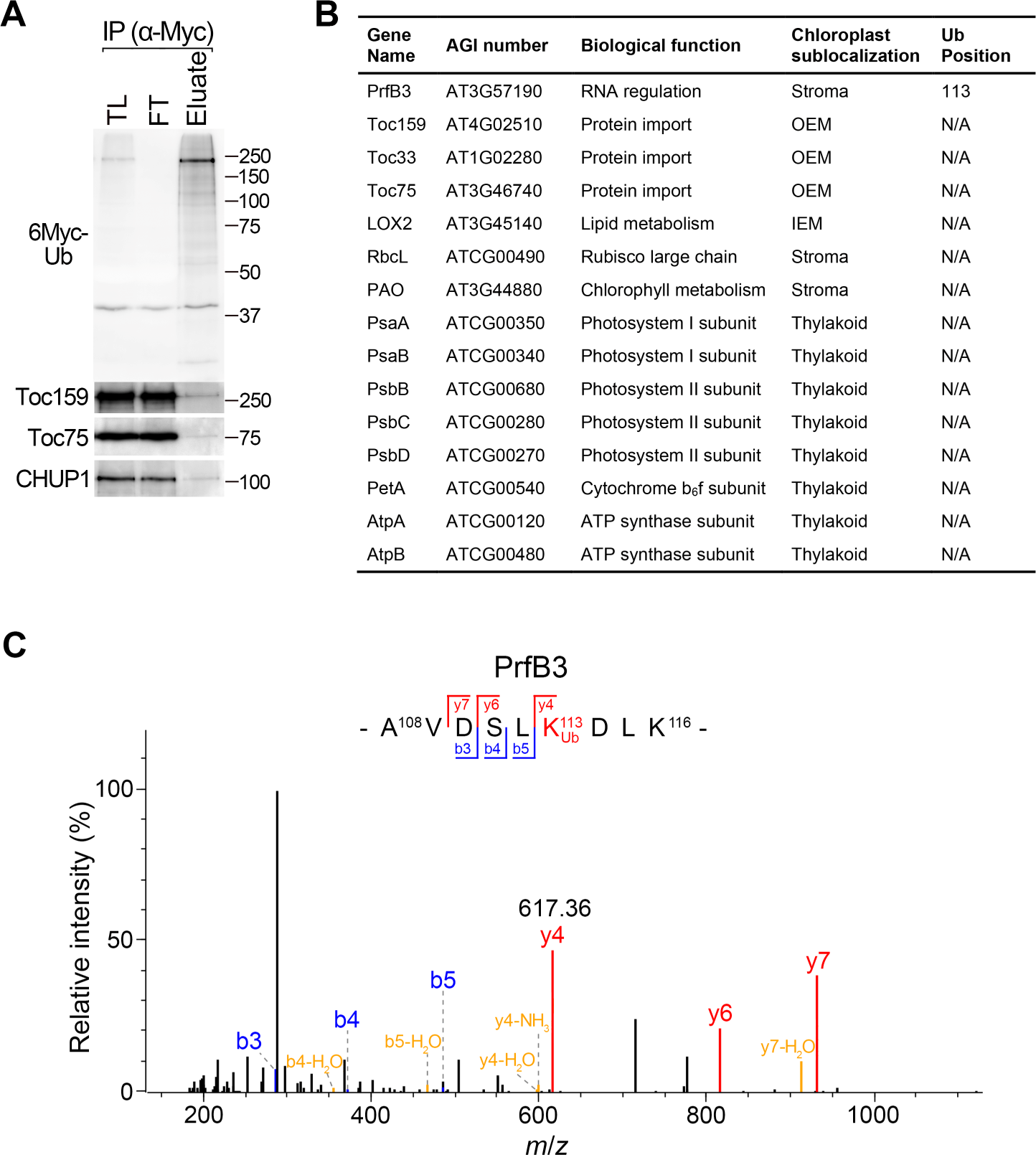
Identification of chloroplast ubiquitination by immunoprecipitation. (**A**) Detection of ubiquitinated proteins in chloroplasts purified from plants expressing Myc-tagged ubiquitin by immunoprecipitation and immunoblotting. Transgenic seedlings expressing ubiquitin with an N-terminal 6×Myc tag (6Myc-Ub) were used to isolate chloroplasts. The 6Myc-Ub chloroplasts were subjected to lysis and membrane solubilization prior to immunoprecipitation (IP) using anti-c-Myc affinity gel. Total lysate (TL; before immunoprecipitation was initiated), IP flow-through (FT), and IP eluate samples were analysed by immunoblotting using anti-Myc and the other indicated antibodies. Ubiquitinated proteins were enriched in the eluted samples, as indicated by the high molecular weight smears, compared with the input sample. (**B** and **C**) Identification of ubiquitinated chloroplast proteins by immunoprecipitation and mass spectrometry. The 6Myc-Ub elution sample shown in A was analysed by LC-MS/MS, and the data were analysed by MaxQuant. This identified six peptides for ubiquitin (covering 78.4% of the whole ubiquitin protein sequence), as well as the chloroplast proteins listed (**B**) (OEM, outer envelope membrane; IEM, inner envelope membrane). After trypsin digestion, a di-glycine remnant of ubiquitin remains covalently linked to target lysine residues of modified proteins (K-ε-GG), and this is identified by an increased mass of 114 kDa of the lysine residue. A ubiquitinated peptide was detected for the stromal protein PrfB3 (B,C), and the associated fragmentation spectrum is shown (C). The y and b ions are shown in red and blue, respectively, while ions with H_2_O or NH_3_ shift are shown in orange. The b ions appear to extend from the N-terminus, and the y ions appear to extend from the C-terminus. The corresponding peptide sequence is presented, with the ubiquitinated K residue is marked in red. The *m/z* value of the corresponding fragment is indicated in black.

**Fig. S2.**
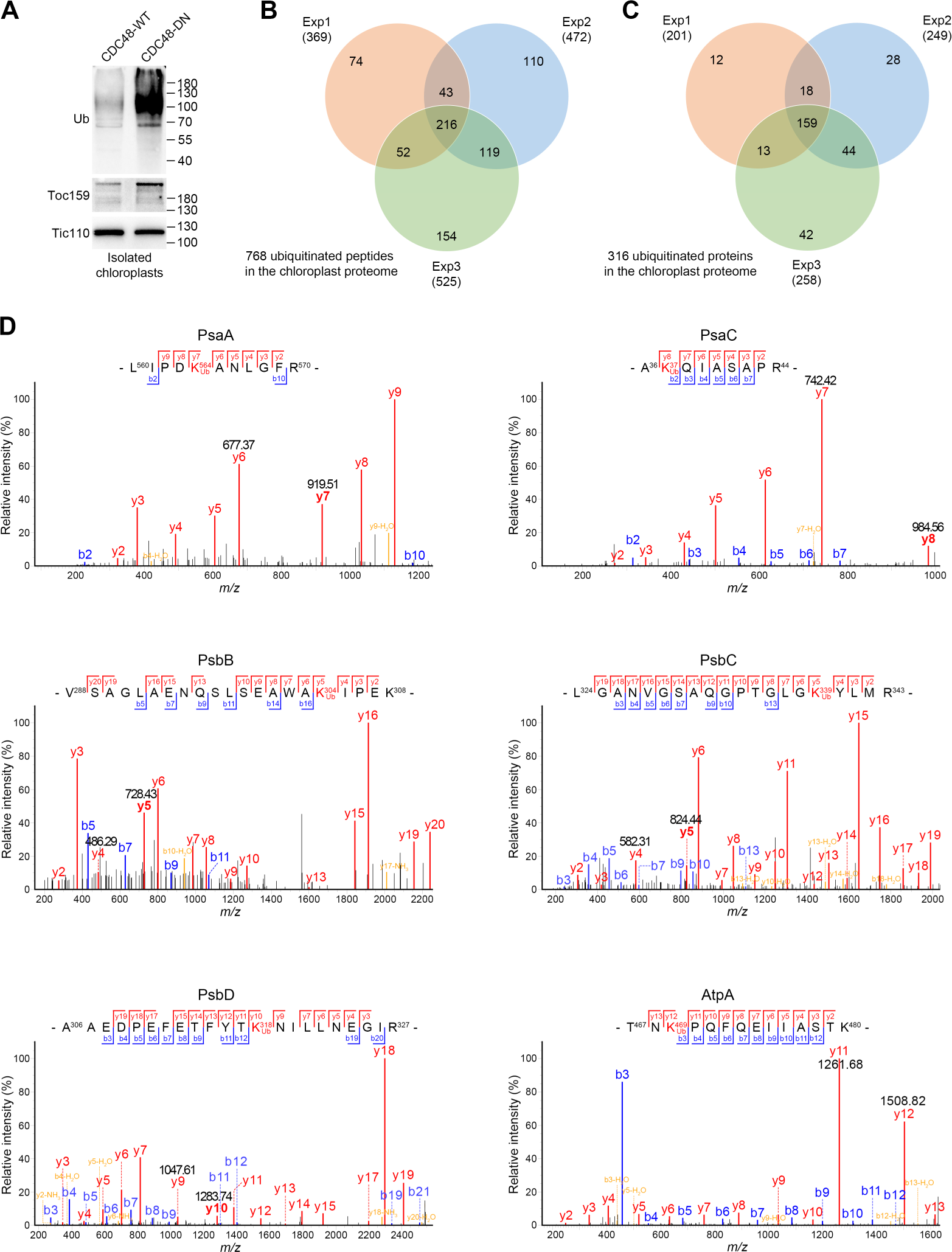
Identification of ubiquitinated chloroplast proteins by di-Gly ubiquitinomics. (**A**) Detection of the accumulation of ubiquitinated proteins in chloroplasts isolated from CDC48-DN plants by immunoblotting. Chloroplasts were isolated from CDC48-WT and CDC48-DN transgenic seedlings that had been induced with estradiol for two days. Induction of CDC48-DN was used to elevate the levels of ubiquitinated chloroplast proteins by blocking the CHLORAD system, in order to increase the chance of capturing ubiquitination events that would otherwise be below the level of detection, due to low stoichiometry or abundance. The purified chloroplasts were analysed by immunoblotting using anti-ubiquitin and the other indicated antibodies. (**B** and **C**) Identification of ubiquitinated proteins and peptides in CDC48-DN chloroplasts by di-Gly ubiquitinomics. Proteins extracted from CDC48-DN chloroplasts like those shown in A were digested with trypsin and subjected to immunoprecipitation analysis using an anti-K-ε-GG antibody. The recovered ubiquitinated peptides were submitted to LC-MS/MS analysis, and the data were analysed by MaxQuant. Three independent biological replicates were analysed (Exp1-3), and the overlaps among the identified ubiquitinated peptides (B) and proteins (C) are shown. (**D**) Identification of ubiquitination sites in several chloroplast-encoded proteins by mass spectrometry. Many of the identified ubiquitinated proteins (B,C) were key photosynthetic complex components, and several of these are encoded by the chloroplast genome: PsaA, PsaC (PSI complex), PsbB, PsbC, PsbD (PSII complex), and AtpA (ATP synthase complex). Peptide sequences and representative spectra are shown for these proteins. The y and b ions are shown in red and blue, respectively, while ions with H_2_O or NH_3_ shift are shown in orange. The b ions appear to extend from the N-terminus, and the y ions appear to extend from the C-terminus. The ubiquitinated K residues are marked in red in the peptide sequences. The *m/z* values of the corresponding fragments are indicated in black.

**Fig. S3.**
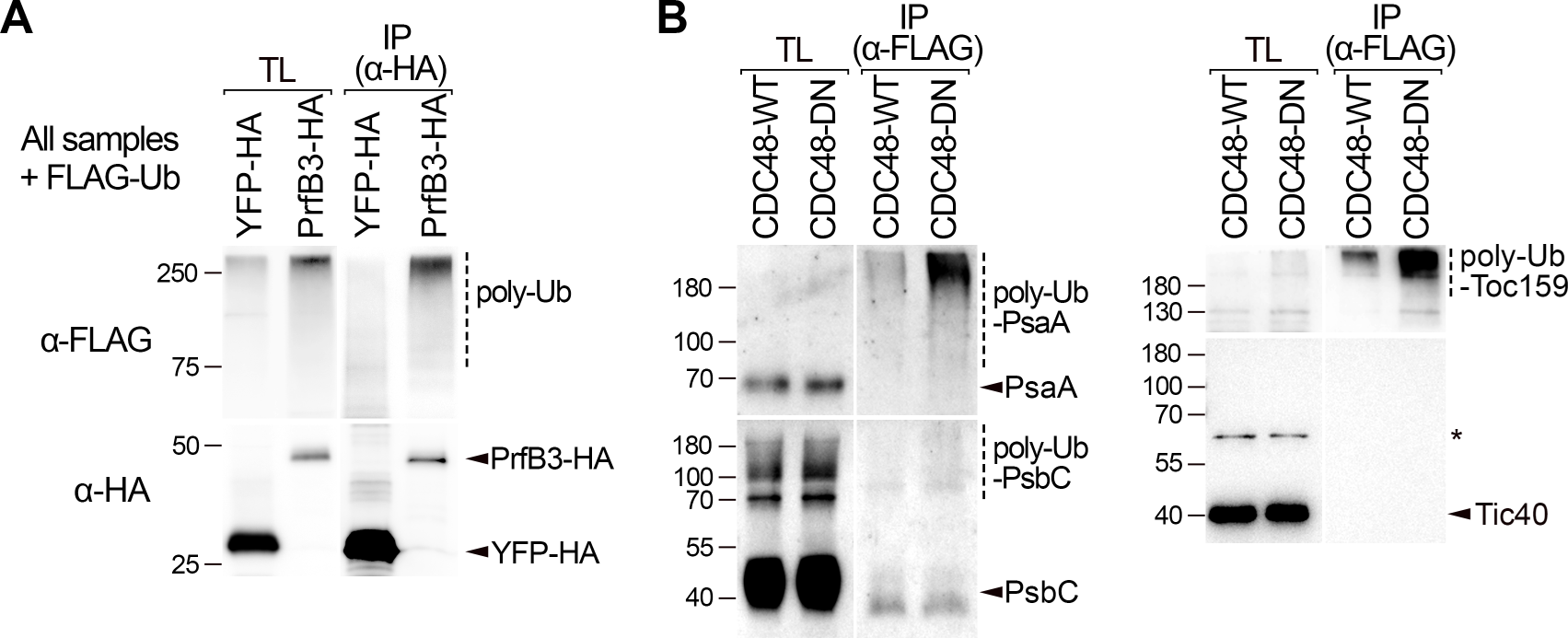
Validation of the ubiquitination of proteins of the chloroplast interior. (**A**) Validation of stromal protein PrfB3 using an in vivo ubiquitination assay. Wild-type protoplasts were cotransformed with constructs encoding HA-tagged PrfB3 (PrfB3-HA) and FLAG-tagged ubiquitin (FLAG-Ub), and the cells were subjected to immunoprecipitation (IP) analysis using anti-HA affinity gel. Total lysate (TL; before IP was initiated) and eluted IP samples were then analysed by immunoblotting using antibodies against: the FLAG tag, to detect polyubiquitinated forms (poly-Ub) of PrfB3-HA or YFP-HA; and the HA tag, to verify that the fusion proteins were present in the samples. Positions of molecular weight markers (sizes in kD) are shown to the left of the images. (**B**) Validation of chloroplast-encoded proteins PsaA and PsbC using an in vivo ubiquitination assay. Protoplasts isolated from CDC48-WT and CDC48-DN transgenic plants were transfected with the FLAG-Ub construct and subjected to estradiol induction. The cells were then analysed by IP using anti-FLAG M2 affinity gel; in this case, ubiquitinated proteins were enriched instead of the target proteins, due to the difficulty of expressing tagged-chloroplast-encoded proteins in plants. Samples (TL, IP) were then analysed by immunoblotting using antibodies against the proteins indicated to the right of the images. Polyubiquitinated forms (poly-Ub) of PsaA and PsbC (and of the known CHLORAD substrate, Toc159) were detected in the IP samples, and these were distinctly larger than their unmodified forms in the TL samples. The poly-Ub forms also increased markedly in abundance in CDC48-DN, suggesting that they are normally subject to CHLORAD degradation. Tic40, which is not a CHLORAD substrate, acted as a negative control. The asterisk indicates a non-specific band. Molecular weight markers are indicated as in A.

**Fig. S4.**
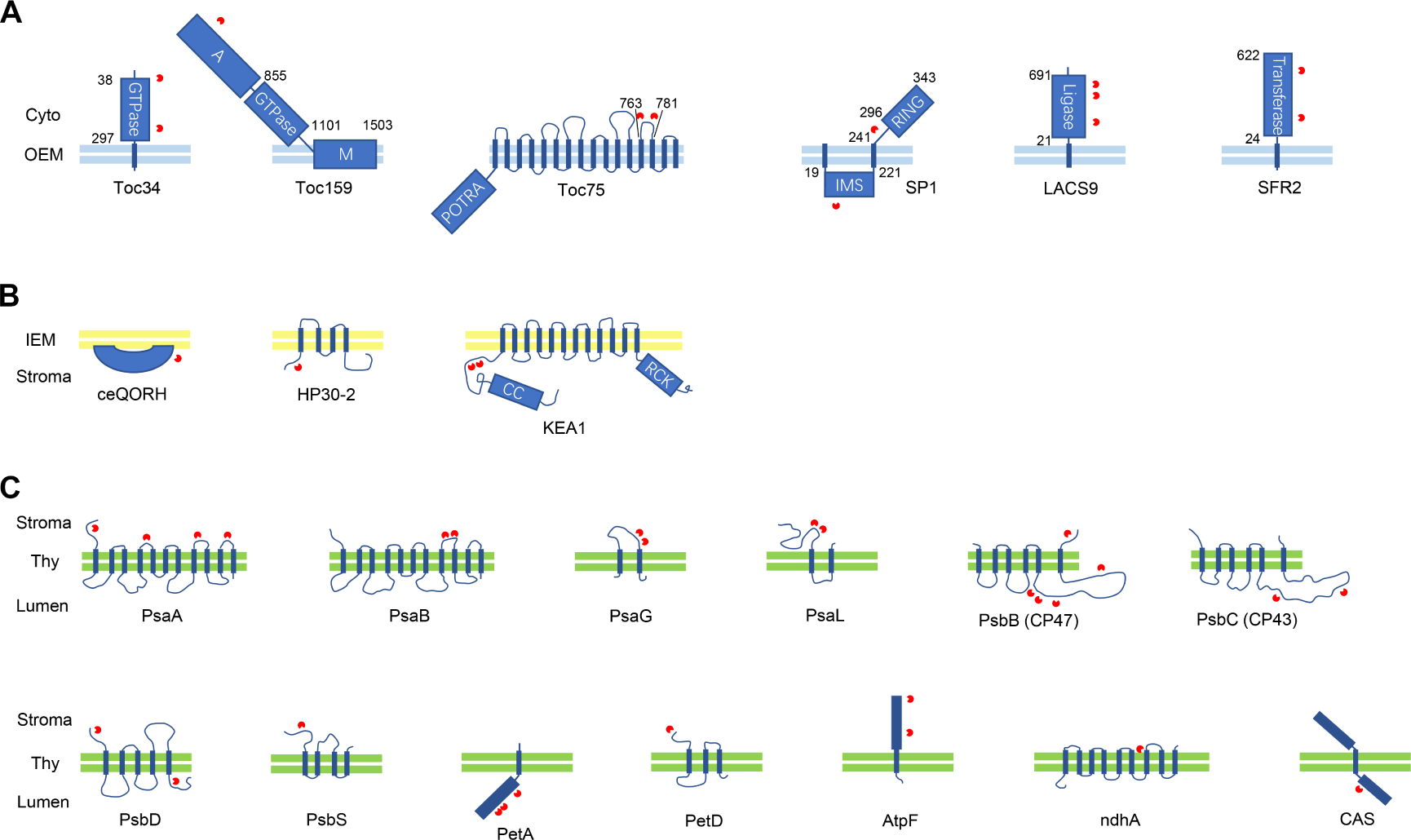
Topological information on the ubiquitination sites of chloroplast proteins. Schematic diagrams are shown for a representative selection of ubiquitinated chloroplast membrane proteins. The corresponding dataset is shown in table S3. The indicated proteins are resident in the outer envelope membrane (OEM; A), the inner envelope membrane (IEM; B), and the thylakoid membrane (Thy; C). Red symbols indicate the positions of ubiquitinated K residues. Numbers indicate the amino acid positions of protein domains; known domains are represented as dark blue shapes. Coloured double lines (light blue, yellow, green) indicate the relevant chloroplast membrane lipid bilayers. Cyto, cytosol. The noticeable lack of ubiquitination in the transmembrane domains implies that the ubiquitination of CHLORAD substrates occurs in situ and is domain specific; alternatively, it might reflect a technical difficulty with the identification of hydrophobic peptides by LC-MS/MS.

**Fig. S5.**
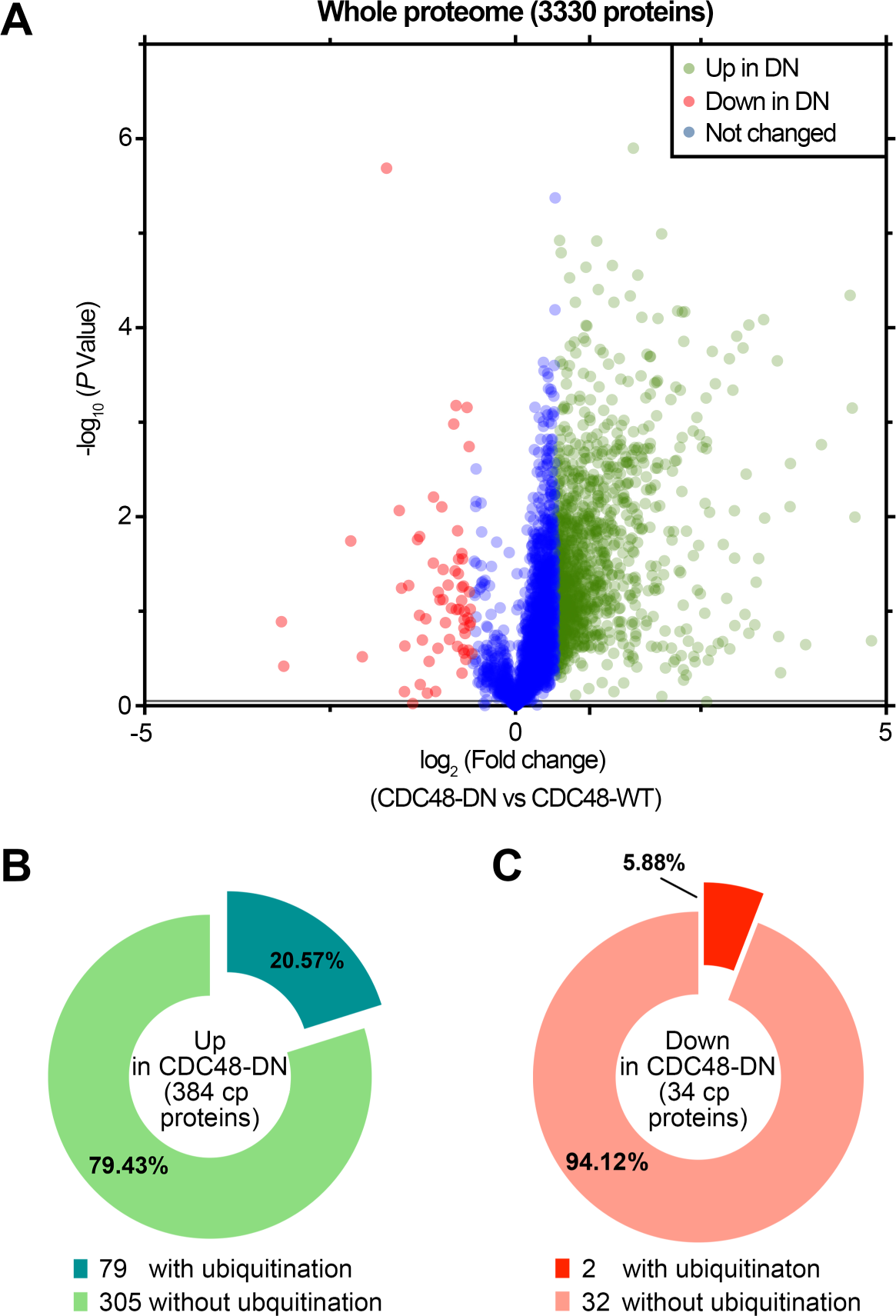
Additional information on the quantitative proteomic analysis. (**A**) Volcano plot representation of the data for all proteins identified in the quantitative proteomic analysis comparing CDC48-DN and CDC48-WT plants, before filtering for chloroplast-related proteins (Fig. 3). The graph shows -log_10_ *P* values plotted against log2 fold changes; proteins deemed to be showing a significant difference in abundance in CDC48-DN plants, relative to the CDC48-WT control, are indicated in green (Up) or red (Down). The corresponding dataset is show in table S5. (**B** and **C**) Pie charts showing the proportion of chloroplast proteins identified in the quantitative proteomics analysis with at least one ubiquitination site. Chloroplast (cp) proteins over-accumulated in CDC48-DN (B), or reduced in CDC48-DN (C), were compared with the chloroplast ubiquitinome (Fig. 2), and in each case the percentage of proteins with detected ubiquitination is shown. The results show that proteins accumulated in CDC48-DN chloroplasts are more prone to be ubiquitinated, which suggests that chloroplast proteins are degraded by the UPS, via CDC48.

**Fig. S6.**
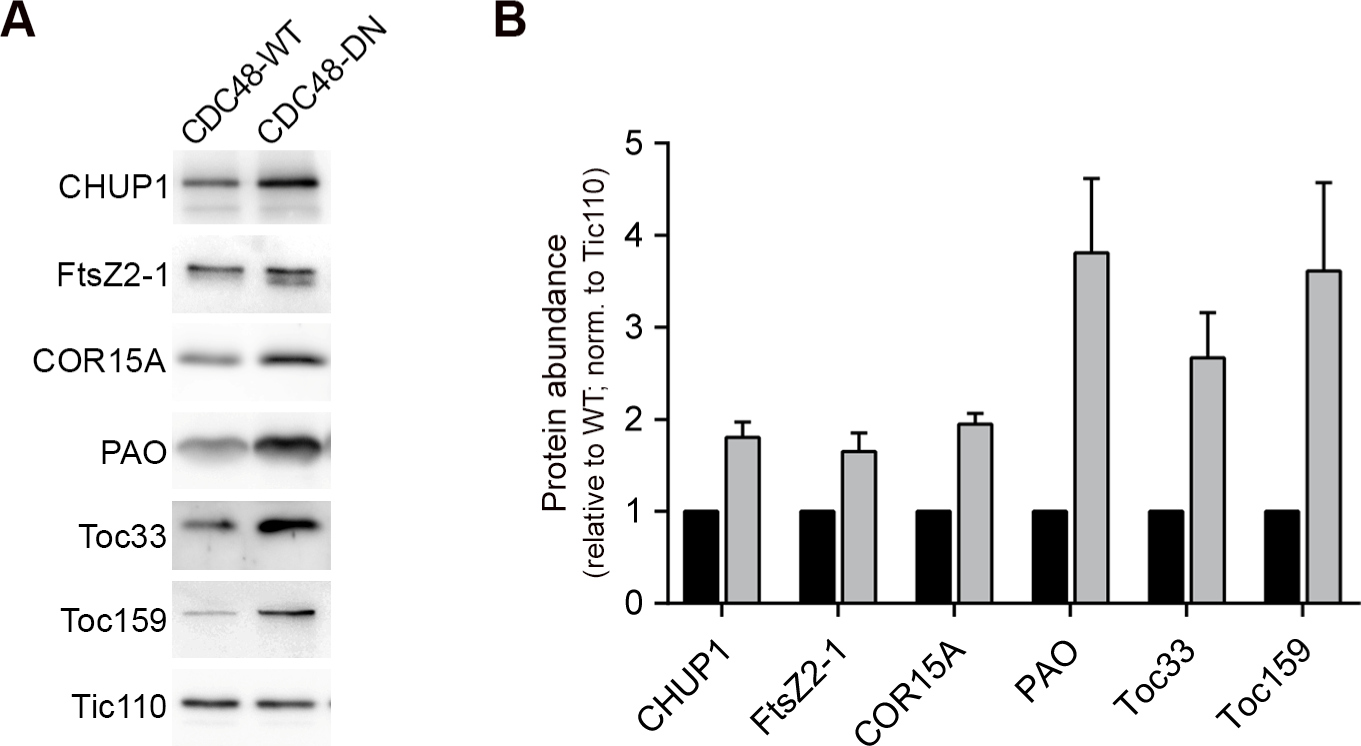
Validation of the quantitative proteomics by using native protein antibodies. Transgenic CDC48-WT and CDC48-DN plants were induced with estradiol for two days, and total protein extracts from these plants were analysed by immunoblotting. The antibodies used were against a selection of chloroplast proteins that showed over-accumulation in the quantitative proteomics analysis (Fig. 3); i.e., proteins deemed to be candidate CHLORAD substrates. The selection of proteins for analysis here was based in part on the availability of antibodies against the native protein. The proteins analysed also had different sub-organellar localizations, as follows: CHUP1 (OEM), FtsZ2-1 (IEM), COR15A, PAO (both stroma). Toc33 and Toc159, which are known CHLORAD substrates, were used as positive controls; Tic110, which is not a CHLORAD substrate, was used as a sample normalization control. Typical immunoblotting results are shown (A). Band intensities for the candidate substrates were quantified and normalized to equivalent data for Tic110 (B). Data are means ± SEM from three biological replicates. The data show that the selected proteins specifically over-accumulate in CDC48-DN plants, verifying the proteomics data and suggesting that they are bona fide CHLORAD substrates.

**Fig. S7.**
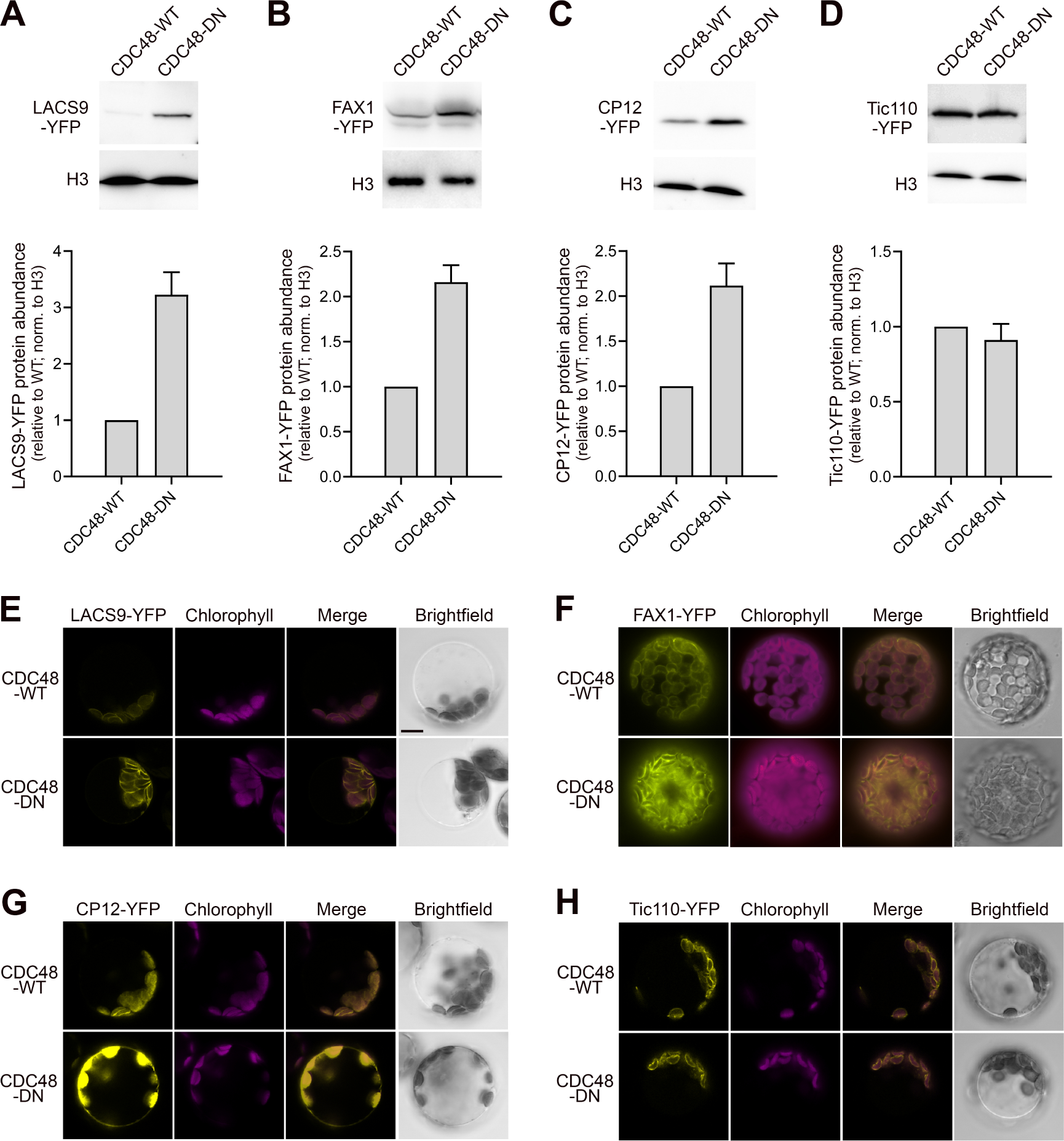
Validation of the quantitative proteomics by analysing YFP-tagged proteins. (**A** to **D**) Verifying the effect of CDC48-DN expression on candidate CHLORAD substrates by immunoblotting. For proteins that showed over-accumulation in the quantitative proteomics analysis (Fig. 3), but with no available antibody against the native protein, a tagging approach was employed for validation. Protoplasts from CDC48-WT and CDC48-DN transgenic plants were transiently transformed with constructs encoding fusion proteins with C-terminal YFP tags. The candidate substrate proteins analysed had different sub-organellar localizations, as follows: LACS9 (OEM; A), FAX1 (IEM; B), CP12 (stroma; C), Tic110 (control; D). The transfected protoplasts were incubated with estradiol for 15-18 h to induce CDC48 transgene expression, and then analysed by immunoblotting using antibodies against: the YFP tag, to detect the fusion proteins; and H3, as an internal sample normalization control. Typical immunoblotting results are shown (upper panels). Band intensities in the experiment shown, and in two additional similar experiments, were quantified; the values obtained for the YFP fusions were normalized using corresponding H3 values. Data are means ± SEM from three different experiments. The data show that the fusion proteins specifically accumulate in CDC48-DN protoplasts, verifying the proteomics data and suggesting that the analysed candidates are bona fide CHLORAD substrates. (**E** to **H**) Verifying the effect of CDC48-DN expression on candidate CHLORAD substrates by fluorescence microscopy. Protoplasts from CDC48-WT and CDC48-DN transgenic plants expressing the YFP-tagged proteins described in A-D were analysed by confocal microscopy. Chlorophyll autofluorescence was used to determine the localization of the YFP fluorescence signals relative to the chloroplasts. The LACS9-YFP (E), FAX1-YFP (F), and Tic110-YFP (H) signals all showed localization to the chloroplast envelopes, whereas the CP12-YFP signal (G) showed localization inside the chloroplasts; thus, all fusion proteins displayed the correct localization pattern. The intensity of the YFP signals for LACS9-YFP, FAX1-YFP and CP12-YFP increased markedly in CDC48-DN cells, implying that these proteins are degraded by CHLORAD at the chloroplasts. Tic110 is not a CHLORAD substrate, and so the intensity of the Tic110-YFP signal was unchanged in CDC48-DN cells. Brightfield images confirmed the intactness of the protoplasts. Scale bar, 10 μm. These data support a role for CHLORAD in degrading proteins resident in the chloroplast interior, rather than un-imported chloroplast preproteins that may exist in the cytosol.

**Fig. S8.**
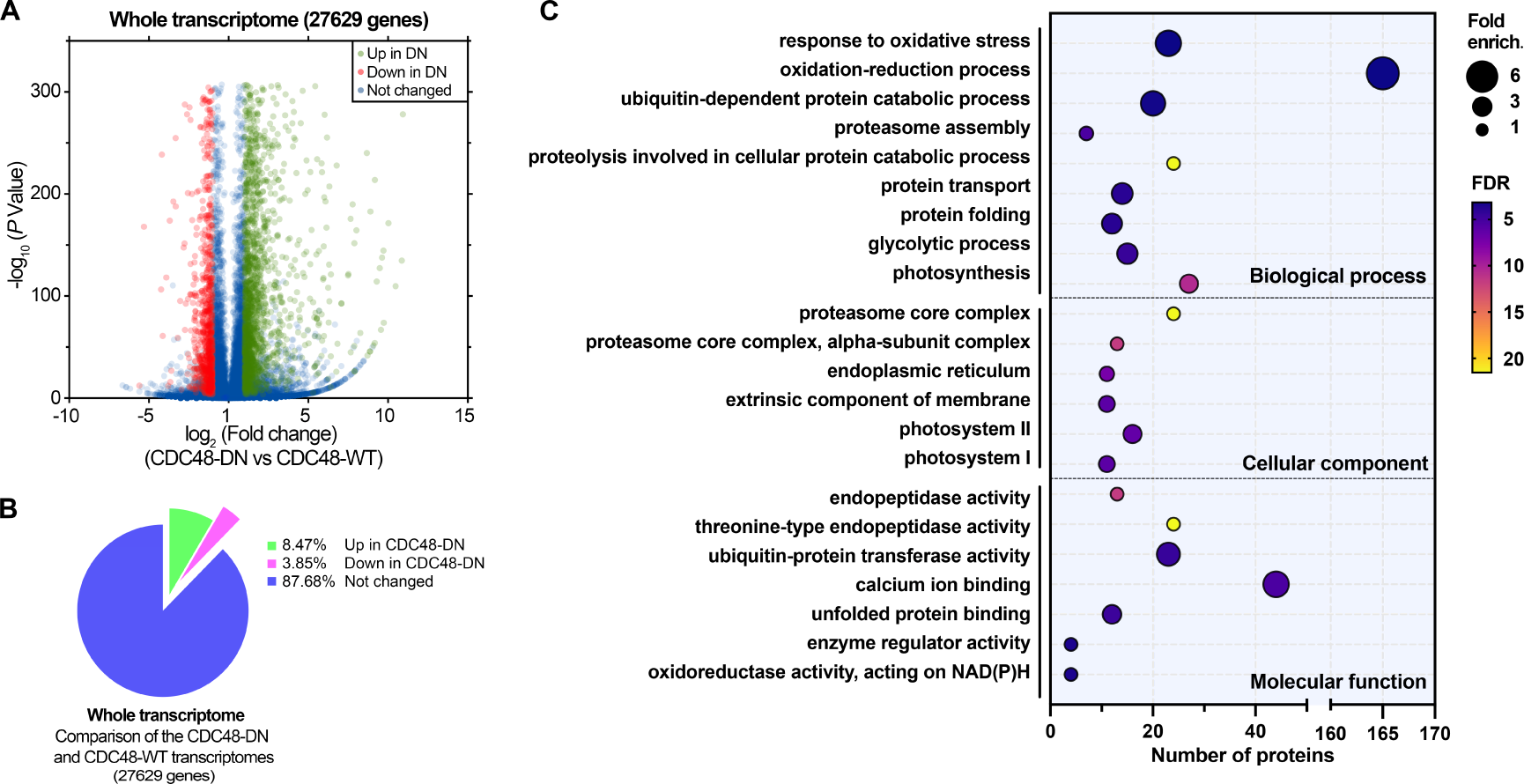
Additional information on the RNA-seq transcriptomic analysis. (**A**) Volcano plot representation of the data for all genes identified in quantitative transcriptomic analysis comparing CDC48-DN and CDC48-WT plants, before filtering for chloroplast-related genes (Fig. 3). The graph shows -log_10_ *P* values plotted against log2 fold changes; genes deemed to be showing a significant difference in expression in CDC48-DN plants, relative to the CDC48-WT control, are indicated in green (Up) or red (Down). The corresponding dataset is show in table S8. (**B**) Pie chart showing the proportion of mRNAs in the whole transcriptome that are differentially expressed in CDC48-DN plants, as determined by the quantitative transcriptomic analysis. In general, more genes were up-regulated than down-regulated in response to CDC48-DN expression. In contrast, chloroplast-related genes showed the opposite trend, with more genes found to be down-regulated (Fig. 3D). (**C**) Dot plot showing significantly overrepresented GO terms for mRNAs that are differentially expressed in CDC48-DN plants. Dot size indicates overrepresentation (fold enrichment) compared to the whole genome. Dot colour indicates False Discovery Rate (FDR; -log_10_ [*P* value]), where higher FDR values indicate more statistically significant enrichment. Dots are not shown for terms lacking statistically significant (*P* < 0.05) enrichment. In general, ubiquitin-dependent proteolytic processes are upregulated, whereas photosynthesis-related pathways are down-regulated.

**Fig. S9.**
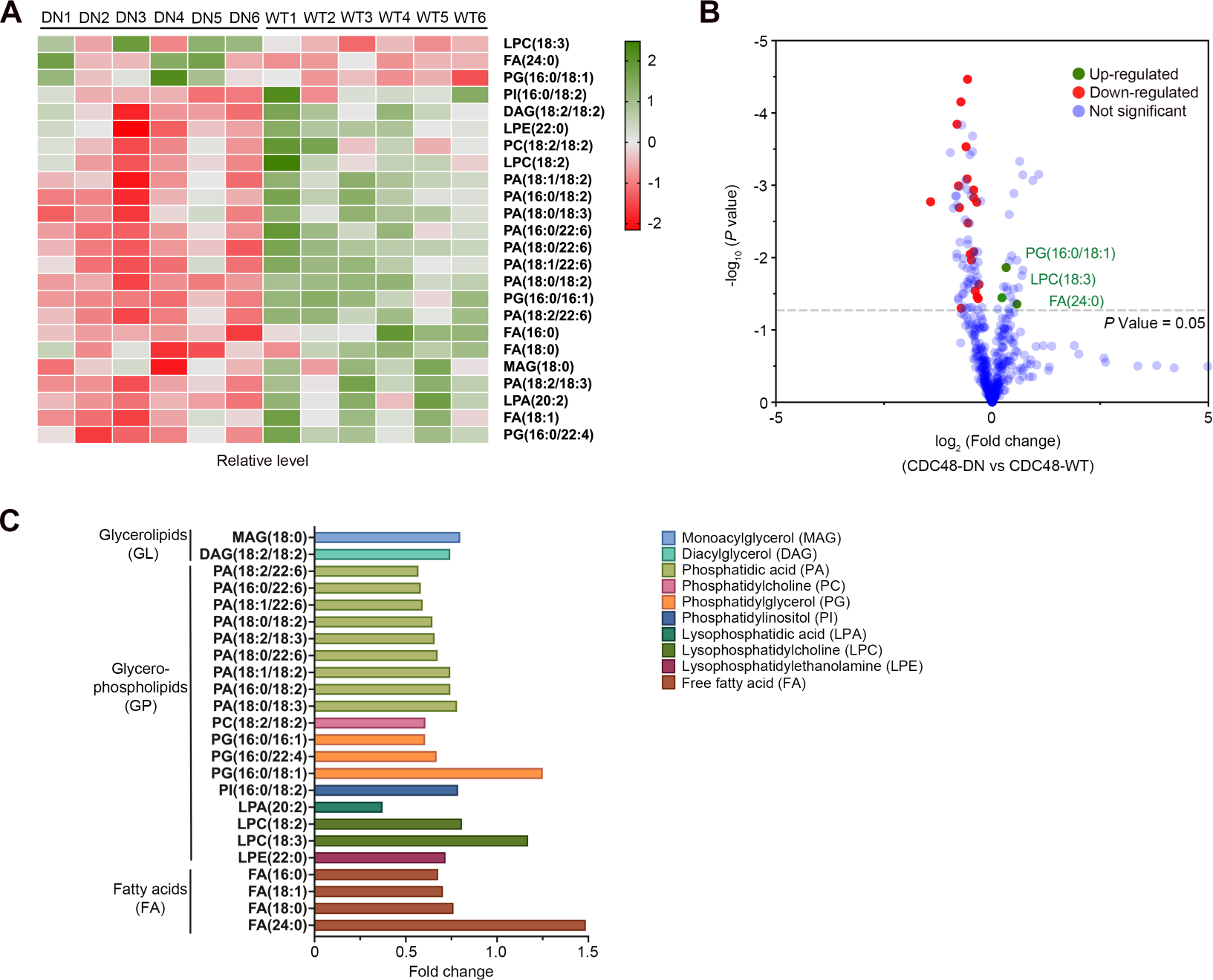
Additional information on the fatty acid/lipid measurements. (**A**) Heatmap showing the distinct FA and lipid profiles of CDC48-WT and CDC48-DN plants. Heatmap was applied to display the relative amounts of FAs and lipids, with a green colour gradient indicating high amounts and a red colour gradient indicating low amounts. Those FA and lipid species with significant changes in the CDC48-WT and CDC48-DN plant groups (six biological replicates for each group) were analysed in the heatmap (VIP [Variable Importance in Projection] > 1, *P* < 0.05). The numbers in the FA/lipid names describe the relevant FA chains; these numbers are presented in the format (number of carbons in the FA chain) : (number of double bonds in the FA chain). (**B**) Volcano plot representation of the data for all FAs and lipids profiled. The graph shows -log_10_ *P* values plotted against log2 fold changes; species deemed to be showing a significant difference in abundance in CDC48-DN plants, relative to the CDC48-WT control, are indicated in green (Up) or red (Down) (VIP > 1, P < 0.05). The corresponding dataset is shown in table S9. (**C**) Quantification of the changes in specific FA and lipid species in CDC48-DN. Bar graph showing relative amounts (fold change values) of species that were significantly changed in CDC48-DN plants, relative to CDC48-WT plants. The species shown are the same as those presented in A. Full names of the FA/lipid species are indicated to the right.

